# CDCA7 is a hemimethylated DNA adaptor for the nucleosome remodeler HELLS

**DOI:** 10.1101/2023.12.19.572350

**Authors:** Isabel E. Wassing, Atsuya Nishiyama, Moeri Hiruta, Qingyuan Jia, Reia Shikimachi, Amika Kikuchi, Keita Sugimura, Xin Hong, Yoshie Chiba, Junhui Peng, Christopher Jenness, Makoto Nakanishi, Li Zhao, Kyohei Arita, Hironori Funabiki

## Abstract

Mutations of the SNF2 family ATPase HELLS and its activator CDCA7 cause immunodeficiency-centromeric instability-facial anomalies (ICF) syndrome, characterized by hypomethylation at heterochromatin. The unique zinc-finger domain, zf-4CXXC_R1, of CDCA7 is widely conserved across eukaryotes but is absent from species that lack HELLS and DNA methyltransferases, implying its specialized relation with methylated DNA. Here we demonstrate that zf-4CXXC_R1 acts as a hemimethylated DNA sensor. The zf-4CXXC_R1 domain of CDCA7 selectively binds to DNA with a hemimethylated CpG, but not unmethylated or fully methylated CpG, and ICF disease mutations eliminated this binding. CDCA7 and HELLS interact via their N-terminal alpha helices, through which HELLS is recruited to hemimethylated DNA. While placement of a hemimethylated CpG within the nucleosome core particle can hinder its recognition by CDCA7, cryo-EM structure analysis of the CDCA7-nucleosome complex suggests that zf-4CXXC_R1 recognizes a hemimethylated CpG in the major groove at linker DNA. Our study provides insights into how the CDCA7-HELLS nucleosome remodeling complex uniquely assists maintenance DNA methylation.

## Introduction

DNA methylation is a broadly observed epigenetic modification in living systems, playing diverse functions in transcriptional regulation, transposable element silencing, as well as innate immunity (*1–4*). As genomic DNA methylation profiles dynamically change during development, aging, and evolution, alterations in DNA methylation patterns are linked to transgenerational epigenetic changes, speciation, and diseases such as cancers and immunodeficiency (*5–9*). One such disease is immunodeficiency–<u>c</u>entromeric instability–<u>f</u>acial anomalies (ICF) syndrome. ICF patient cells exhibit hypomethylation of heterochromatic regions, particularly at the juxta-centromeric heterochromatin of chromosome 1 and 16 (*10, 11*). Mutations in four genes are known to cause ICF syndrome; the *de novo* DNA methyltransferase DNMT3B, the SNF2-family ATPase HELLS (also known as LSH, SMARCA6 or PASG), the HELLS activator CDCA7, and the transcription factor ZBTB24, which is critical for the expression of CDCA7 (*12–16*). In addition, compound mutations of UHRF1, a critical regulator of maintenance DNA methylation, cause atypical ICF syndrome (*17*), supporting a further causal relationship between defective DNA methylation and the disease. The importance of HELLS and its plant ortholog DDM1 in DNA methylation has been established in vertebrates and in plants (*18–25*), and it has been suggested that the nucleosome remodeling activity of HELLS/DDM1 facilitates DNA methylation (*26, 27*). However, it remains unclear why a role in promoting DNA methylation is uniquely carried out by HELLS/DDM1 among several other coexisting SNF2-family ATPases with similar nucleosome remodeling activity, such as SNF2 (SMARCA2/4), INO80, and ISWI (SMARCA1/5) (*28*).

In eukaryotes, DNA methylation is primarily observed as 5-methylcytosine (5mC), commonly in the context of CpG sequences, where both cytosines in the complementary DNA strands are symmetrically (i.e., fully) methylated. 5mC methylation mechanisms can be functionally classified as maintenance methylation or *de novo* methylation (*29*). Whereas *de novo* methylation, which is commonly mediated by DNMT3-family proteins, does not depend on preexisting 5mC on the template DNA, maintenance methylation, mediated by DNMT1-family proteins, occurs at hemimethylated CpGs, which are generated upon replication of fully methylated DNA. So far, the SRA domain of UHRF1 is the only established eukaryotic protein module that specifically recognizes hemimethylated CpGs (*30–32*). Through its E3 ubiquitin ligase activity, UHRF1 recruits and activates the maintenance DNA methyltransferase DNMT1 (*33–38*). During DNA replication, UHRF1-mediated dual mono-ubiquitylation of the PCNA-associated factor PAF15 promotes DNMT1 activity to support DNA replication-coupled maintenance DNA methylation (*37*). Additionally, when hemimethylated CpGs elude the imperfect replication-coupled maintenance methylation mechanism, DNMT1 can catalyze maintenance methylation far behind the replication fork. It has been suggested that this replication-uncoupled maintenance DNA methylation acts as a backup mechanism, which is most clearly observed in late-replicating/heterochromatin regions and is supported by UHRF1-mediated histone H3 dual mono-ubiquitylation, which activates DNMT1 (*16, 37, 39*). It was also shown that HELLS accelerates replication-uncoupled maintenance DNA methylation at late-replicating regions in HeLa cells (*39*). Furthermore, it has been reported that HELLS can assist the recruitment of UHRF1 and DNMT1 to chromatin and promote H3 ubiquitylation (*25*). While the observed HELLS-UHRF1 interaction may underlie the importance of HELLS in replication-uncoupled maintenance methylation (*25*), it remains unclear how HELLS is effectively recruited to sites of hemimethylation in this process.

The abundance of nucleosomes, which drastically distort the DNA that wraps around the core histone octamer, affects the accessibility/activity of many DNA-binding proteins (*40*), including DNA methyltransferases (*41–45*). The location of hemimethylated DNA within the nucleosome core particle (NCP) also inhibits its detection by the SRA domain of UHRF1 (*46*). *In vivo*, nucleosomal barriers to DNA methylation can be alleviated by the SNF2-family ATPase HELLS in vertebrates and DDM1 in plants (*26*). Although DDM1 can remodel the nucleosome on its own (*47, 48*), we have previously demonstrated that HELLS alone is inactive and must bind CDCA7 to form the <u>C</u>DCA7-<u>H</u>ELLS ICF-<u>r</u>elated nucleosome <u>r</u>emodeling <u>c</u>omplex (CHIRRC), which exerts DNA-dependent ATPase and nucleosome remodeling activities (*27*). In *Xenopus* egg extracts, CDCA7 is critical for recruiting HELLS to chromatin, but not vice versa. HELLS also interacts with CDCA7 in human cells (*49*). The molecular basis of HELLS-CDCA7 interaction and CDCA7-chromatin interaction has not yet been established.

CDCA7 is characterized by its unique zinc-finger domain zf-4CXXC_R1, which is broadly conserved in eukaryotes (*28*) (fig. S1). CDCA7 homologs with the prototypical zf-4CXXC_R1 domain, containing eleven highly conserved signature cysteine residues and three ICF disease-associated residues, are almost exclusively identified in species that also harbor HELLS/DDM1 and maintenance DNA methyltransferases (DNMT1/MET1 or DNMT5), whereas CDCA7 is almost always lost in species that lack detectable genomic 5mC, such as *Drosophila*, *Tribolium*, *Microplitis*, *Caenorhabditis*, *Schizosaccharomyces*, and *Saccharomyces* (*28*). This coevolution analysis suggests that zf-4CXXC_R1 domain became readily dispensable in species that lack methylated DNA (*28*). However, the function of zf-4CXXC_R1 remains to be defined. Here, we demonstrate that the zf-4CXXC_R1 domain of CDCA7 is a sensor for hemimethylated DNA. Our results help explain how CDCA7 could confer the unique role of HELLS in maintenance DNA methylation.

## Results

### Inhibiting maintenance DNA methylation enriches HELLS and CDCA7 on chromatin

Although CDCA7 coevolved with HELLS and the maintenance DNA methyltransferases (*28*), their mechanistic link remained unclear. The first hint emerged when we observed that HELLS preferentially accumulated on sperm chromatin after incubation in interphase DNMT1-depleted *Xenopus* egg extracts (ΔDNMT1) (Fig. 1A). Adding sperm nuclei to egg extracts promotes functional nuclear formation, upon which DNA replication is rapidly executed between 30-60 min after incubation (*50*). DNA synthesis on the highly methylated sperm chromosomal DNA transiently generates hemimethylated DNA, which immediately induces maintenance DNA methylation by UHRF1 and DNMT1 (*35, 37, 38*). Therefore, when maintenance methylation is inhibited, hemimethylated DNA is expected to accumulate during DNA replication. Indeed, the accumulation of higher molecular weight H3 species, characteristic for mono- and di-ubiquitylated H3, in the DNMT1-depleted extract is in line with the absence of maintenance methylation (Fig. 1A). We thus speculated that the observed enhanced enrichment of HELLS on chromatin in ΔDNMT1 extracts was caused by the accumulation of hemimethylated DNA. Alternatively, as it has been reported that UHRF1 and HELLS interact (*25*), this HELLS enrichment on ΔDNMT1 extract could be caused by chromatin enrichment of UHRF1, which directly binds hemimethylated DNA (*30–32*). To distinguish between these possibilities, we used recombinant mouse DPPA3 (mDPPA3), which binds to UHRF1 and inhibits its association with chromatin (*51–53*). In control egg extracts, DNMT1, UHRF1, HELLS and CDCA7e (a sole CDCA7 paralog present in *Xenopus* eggs) transiently associated with chromatin in S phase (40-60 min after sperm nucleus addition to egg extracts) (Fig. 1B). In the presence of mDPPA3, DNMT1 and UHRF1 failed to associate with chromatin, while CDCA7e and HELLS exhibit robust and continuous chromatin accumulation during the time course (Fig. 1B). These results support the idea that CDCA7e and HELLS are enriched on highly hemimethylated chromatin generated upon DNA replication in the absence of active maintenance DNA methylation. Consistent with this idea, chromatin association of CDCA7e and HELLS was suppressed when DNA replication was inhibited by geminin (fig. S2)(*54*).

**Fig. 1.**
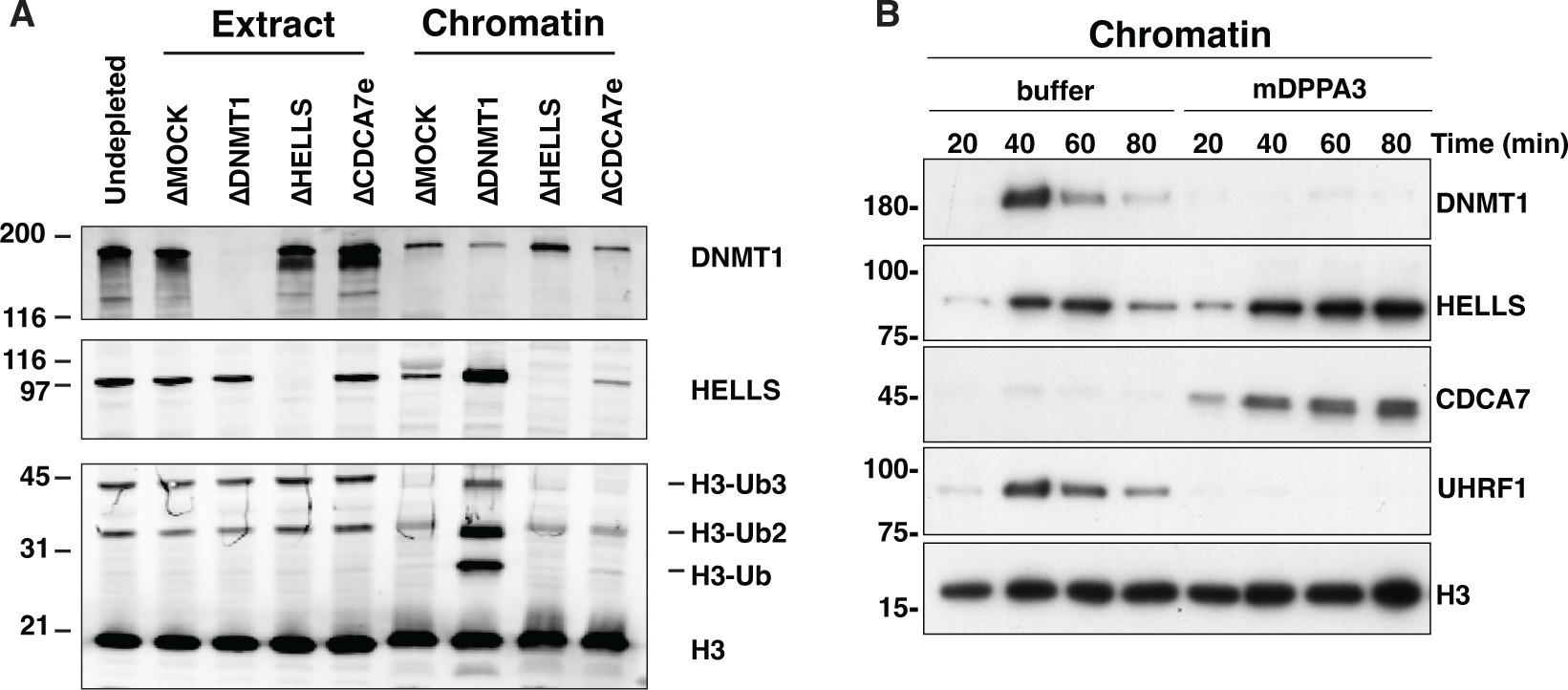
CDCA7 and HELLS accumulate on chromatin upon inhibition of maintenance DNA methylation. **(A)** *X. laevis* sperm nuclei were incubated with interphase egg extracts depleted with mock IgG (ΔMOCK), anti-DNMT1 (ΔDNMT1), anti-HELLS (ΔHELLS), or anti-CDCA7e (ΔCDCA7e) antibodies for 3 h in the presence of cycloheximide. Chromatin was isolated and analyzed by western blotting. **(B)** *X. laevis* sperm nuclei were isolated at indicated time points after incubation with interphase *Xenopus* egg extracts in the presence or absence of 0.5 µM mouse DPPA3 (mDPPA3), a protein that inhibits binding of UHRF1 and DNMT1 to chromatin. Chromatin-associated proteins were analyzed by western blotting.

### CDCA7 zf-4CXXC_R1 domain selectively binds hemimethylated DNA

CDCA7-family proteins are defined by the presence of the unique zf-4CXXC_R1 domain, in which all three identified ICF-disease associated residues are highly conserved (fig. S1) (*28*). Since CDCA7e recruits HELLS to chromatin in *Xenopus* egg extracts but not vice versa (*27*), we explored a possibility that CDCA7e directly recognizes hemimethylated DNA via the zf-4CXXC_R1 domain. To test this hypothesis, beads coupled with unmethylated, hemimethylated, or fully methylated DNA at CpG sites were incubated with *Xenopus* egg extracts. As expected, UHRF1 and ubiquitylated H3 were preferentially enriched on hemimethylated DNA beads (Fig. 2A). Strikingly, CDCA7e was markedly enriched on hemimethylated DNA over unmethylated or fully methylated DNAs (Fig. 2A). When ^35^S-labeled *X. laevis* CDCA7e produced in reticulocyte lysates was assessed for its DNA binding *in vitro*, wildtype CDCA7e but not CDCA7e with any of the ICF disease-associated mutations (R232H, G252V, or R262H) selectively associated with hemimethylated DNA (Fig. 2B, Table S1). Direct and specific binding of CDCA7e to hemimethylated DNA was further confirmed by electrophoretic mobility shift assay using purified recombinant protein and double-stranded oligo-DNA containing a single hemimethylated CpG site (Fig. 2C and D).

**Fig. 2.**
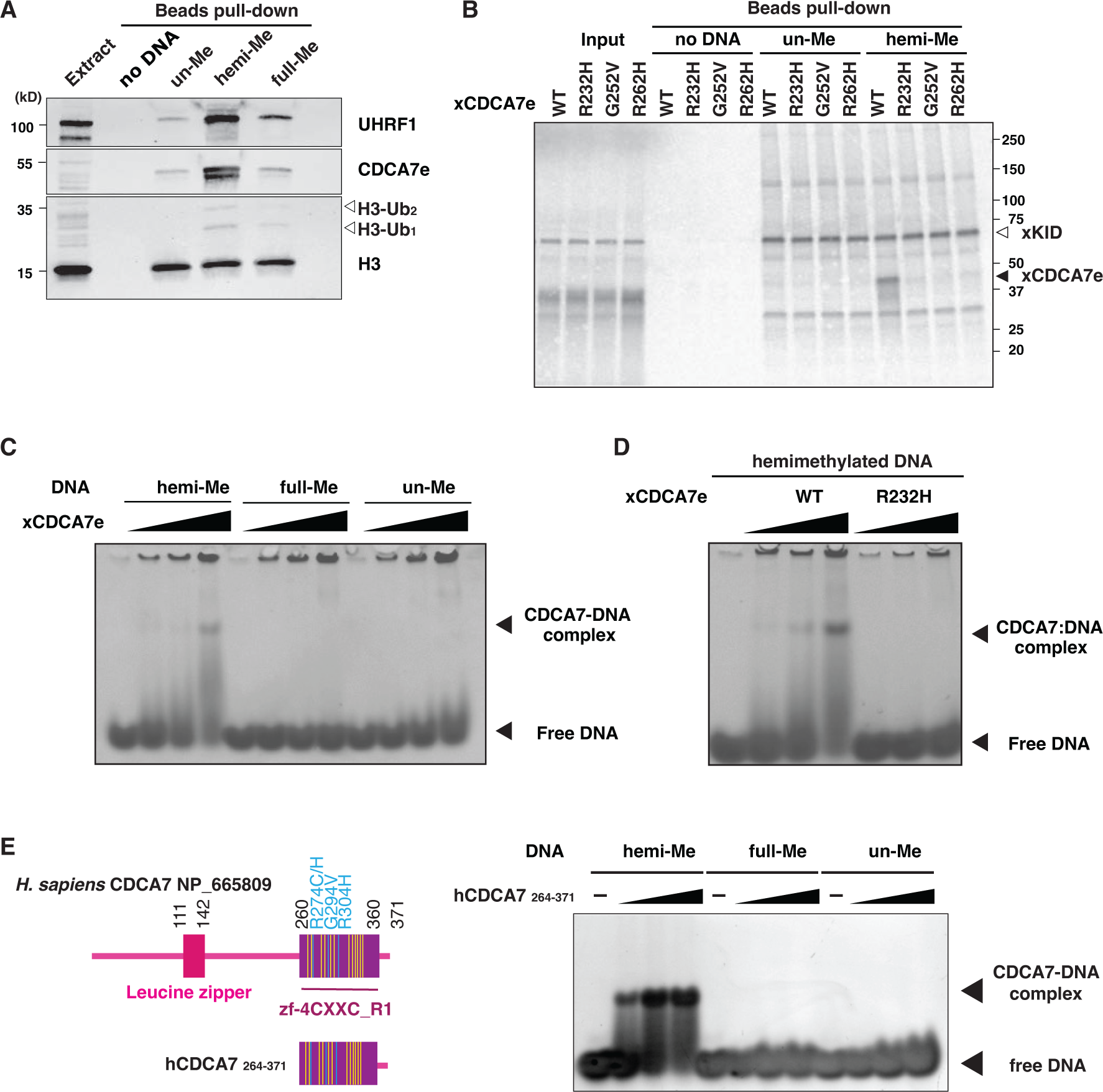
CDCA7 selectively binds hemimethylated DNA. **(A)** Magnetic beads coupled with pBluescript DNA with unmethylated CpGs (un-Me), hemimethylated CpGs (hemi-Me), or fully methylated CpGs (full-Me), were incubated with interphase *Xenopus* egg extracts. Beads were collected after 60 min and analyzed by western blotting. **(B)** ^35^S-labeled *X. laevis* CDCA7e proteins (wildtype or with the indicated ICF3-patient associated mutation) were incubated with control beads, or beads conjugated 200 bp unmethylated or hemimethylated DNA (Table S1). ^35^S-labeled Xkid (*85*), a nonspecific DNA-binding protein, was used as a loading control. Autoradiography of ^35^S-labeled proteins in input and beads fraction is shown. **(C)** and **(D)** Native gel electrophoresis mobility shift assay (EMSA) using recombinant *X. laevis* CDCA7e^WT^ and CDCA7e^R232H^. **(E)** Left: schematic of *H. sapiens* CDCA7 (isoform 2 NP_665809). Positions of the zf-4CXXC_R1 domain (purple), three ICF3-patient mutations (cyan), and conserved cysteine residues (yellow) are shown. Right: EMSA assay using the purified zf-4CXXC_R1 domain (aa 264-371) of *H. sapiens* CDCA7. For C-E, double-stranded DNA oligonucleotides with an unmethylated, hemimethylated or fully-methylated CpG used for protein binding were visualized.

This hemimethylated DNA-specific binding was also observed for human CDCA7. Using the recombinant zf-4CXXC_R1 domain of human CDCA7 (Fig. 2E and fig. S3A), we found that the cysteine-rich segment (aa 264-340 in hCDCA7 NP_665809) of the zf-4CXXC_R1 domain alone does not exhibit any detectable DNA binding capacity (fig. S3B). Adding an N-terminal extension (aa 235-263) to the cysteine-rich segment weakly increased binding to the oligo-DNA with a hemimethylated CpG (fig. S3C). However, extending the cysteine-rich segment to include the evolutionarily conserved C-terminus, which contains two predicted alpha helices, conferred highly selective hemimethylation-dependent DNA binding (Fig. 2E, fig. S1 and S3A). Altogether these results demonstrate that the zf-4CXXC_R1 domain of CDCA7 acts as a hemimethylated DNA-binding module.

### CDCA7 recognizes a hemimethylated CpG at the major groove of linker DNA

Since CDCA7 stimulates nucleosome remodeling activity of HELLS, we asked how the nucleosome could affect recognition of hemimethylated CpG by CDCA7. To address this question by biochemical and structural approaches, we generated the recombinant zf-4CXXC_R1 domain of human CDCA7 (hCDCA7_264-371_ C339S). The C339S substitution was included to improve protein homogeneity during purification while maintaining robust hemimethylated CpG-specific binding (fig. S3D); C339 is not broadly conserved in CDCA7 family proteins and is substituted to serine in *Xenopus* CDCA7e (fig S1). (*28*). Native gel electrophoresis demonstrated that the nucleosome-hCDCA7_264-371_ C339S complex was readily observed when a hemimethylated CpG was positioned at the linker DNA either at its 5’-end or 3’-end (Fig. 3A and table S2). However, the complex formation was undetectable when the hemimethylated CpG was located within the NCP (Fig. 3A).

**Fig. 3:**
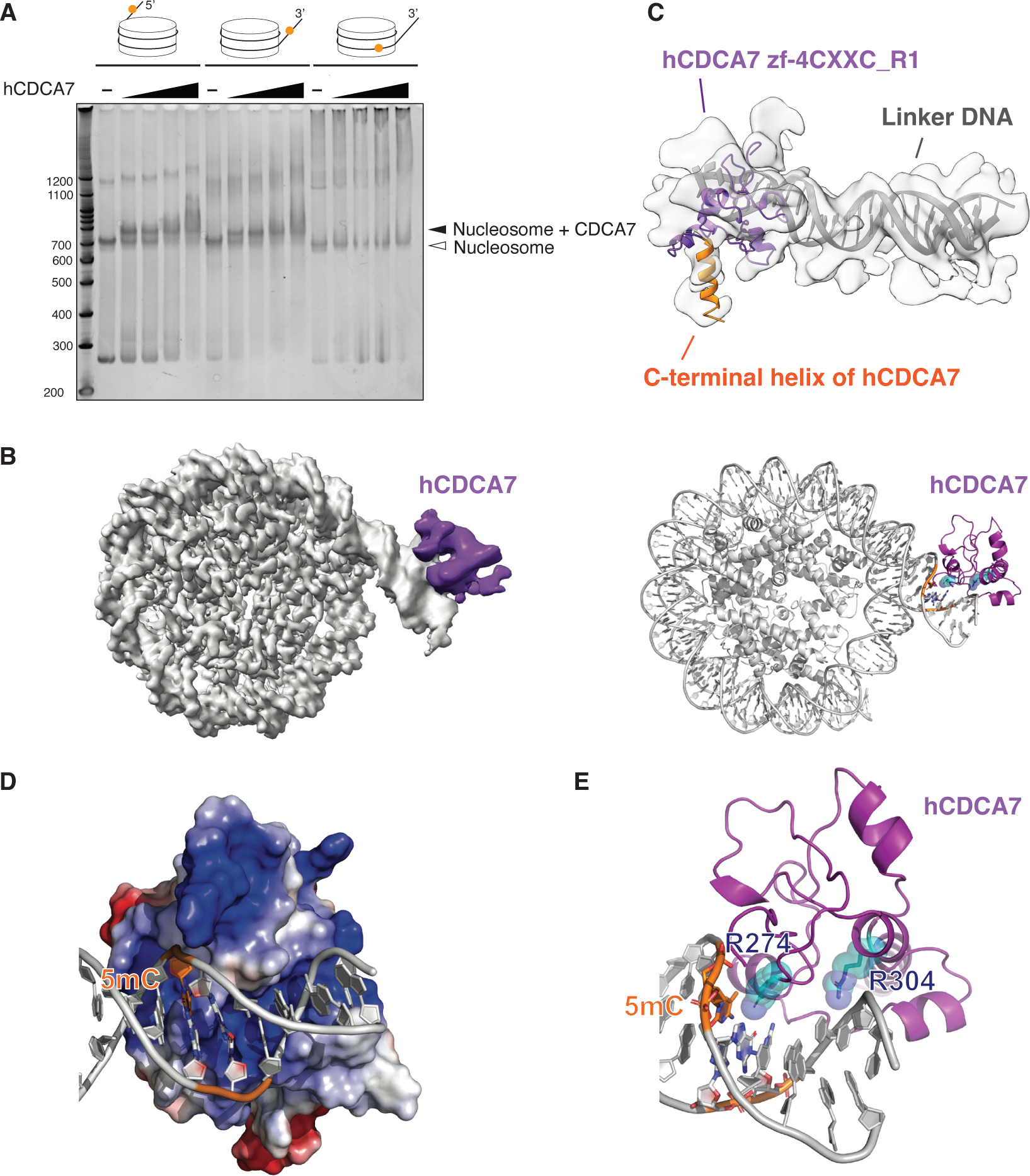
Cryo-EM structure of hCDCA7:nucleosome complex. **(A)** Native gel electrophoresis mobility shift assay analyzing the interaction of hCDCA7_264-371_ C339S with nucleosomes. **(B)** A composite cryo-EM map (left) and the model structure of hCDCA7_264-371_ C339S (generated from AF2) bound to nucleosome harboring a hemimethylated CpG at the 3’-linker DNA (right). The map corresponding to CDCA7 is colored purple. **(C)** Overlay of AF2 model of hCDCA7_264-371_ C339S on the cryo-EM map. **(D)** Electrostatic surface potential of hCDCA7_264-371_, where red and blue indicate negative and positive charges, respectively. Linker DNA is depicted in gray, orange indicates the location of 5-methylcytosine (5mC). **(E)** A model structure of hCDCA7_264-371_ C339S bound to 3’-linker DNA. ICF mutation residues, R274 and R304, are shown as cyan stick model superimposed on a transparent sphere model.

To gain structural insight into CDCA7-hemimethylated DNA interaction, cryogenic electron microscopy (cryo-EM) single particle analysis was conducted on hCDCA7_264-371_ C339S in complex with a mono-nucleosome carrying a hemimethylated CpG at the 3’-linker DNA (fig. S3, fig. S4, S5, Table S2, and Table S3). The initial cryo-EM map showed a density around the major groove of the hemimethylated CpG in the linker DNA, although the density was ambiguous due to the flexibility of the complex (fig. S4). 3D variability analysis and 3D classification generated a cryo-EM map of 3.18 Å resolution for the NCP, where core histones and the phosphate backbone of DNA were clearly resolved, and local refinement and local classification generated a 4.83 Å resolution map for an extra cryo-EM density located outside of the linker DNA (Fig. 3B, fig. S4, S5). This extra density is thought to be hCDCA7_264-371_ bound to linker DNA, as it aligns reasonably well with the AlphaFold2 (AF2)-predicted structure of the zf-4CXXC_R1 domain of human CDCA7 (Fig. 3C) (*55, 56*). First, a notable protrusion of the extra cryo-EM density matches the characteristic C-terminal alpha-helix of hCDCA7 predicted by AF2 (Fig. 3C, orange). Second, fitting the AF2-predicted model of hCDCA7 model structure into the cryo-EM map predicts that the protein surface facing the DNA backbone is positively charged (Fig. 3D). Furthermore, in this structure model, the side chain of R304 and R274, mutated in ICF patients, respectively point toward the DNA backbone and the DNA major groove where the hemimethylated CpG resides (Fig. 3E), consistent with the observed abrogation of hemimethylated CpG binding upon mutating these residues (Fig. 2B, 2D).

### Characterization of the HELLS-CDCA7 interaction interface

Our previous coevolution analysis has shown that the evolutionary preservation of CDCA7 is tightly coupled to the presence of HELLS; while CDCA7 and HELLS were frequently lost from several eukaryote lineages, all the tested eukaryotic species that encode CDCA7 also have HELLS (*28*). As this suggests an evolutionarily conserved function involving both CDCA7 and HELLS, we reasoned that the HELLS-CDCA7 interaction interface is likely also conserved in these species. We employed AF2 structure prediction of HELLS-CDCA7 complex using sequences of HELLS/DDM1 and CDCA7 homologs from diverse eukaryotic species to identify likely CDCA7-HELLS interaction domains (*55, 56*). In all tested cases (*X. laevis* HELLS-CDCA7e, *H. sapiens* HELLS-CDCA7, *H. sapiens* HELLS-CDCA7L, *Ooceraea biroi* (clonal raider ant) HELLS-CDCA7, *Nematostella vectensis* (starlet sea anemone) HELLS-CDCA7, and *Arabidopsis thaliana* DDM1-CDCA7), AF2 predicted the interaction of an N-terminal alpha helix of CDCA7 (aa 74-105 of *X. laevis* CDCA7e) with an N-terminal alpha helix of HELLS/DDM1 (aa 63-96 of *X. laevis* HELLS), as well as multiple segments within the SNF2_N domain of HELLS/DDM1 (Fig. 4A, B and fig. S6). The N-terminal putative CDCA7-binding alpha helix of HELLS corresponds to the previously annotated CC2 (coiled-coil2) segment, while it has been reported that the deletion of the preceding CC1 activates human HELLS by releasing its autoinhibition (*57*). AF2 also predicted an additional shorter CDCA7-binding interface in *X. laevis* and *H. sapiens* HELLS (aa 163-172 in *X. laevis* HELLS) (Fig. 4A, B and fig. S6A-D). The putative interacting alpha helices of CDCA7 and HELLS/DDM1 are evolutionarily conserved in divergent green plant and animal species (Fig. 4C, D, fig. S6C-G and fig. S7), whereas sequence conservation of the second CDCA7-binding interface in HELLS is less clear (Fig. 4E).

**Fig. 4.**
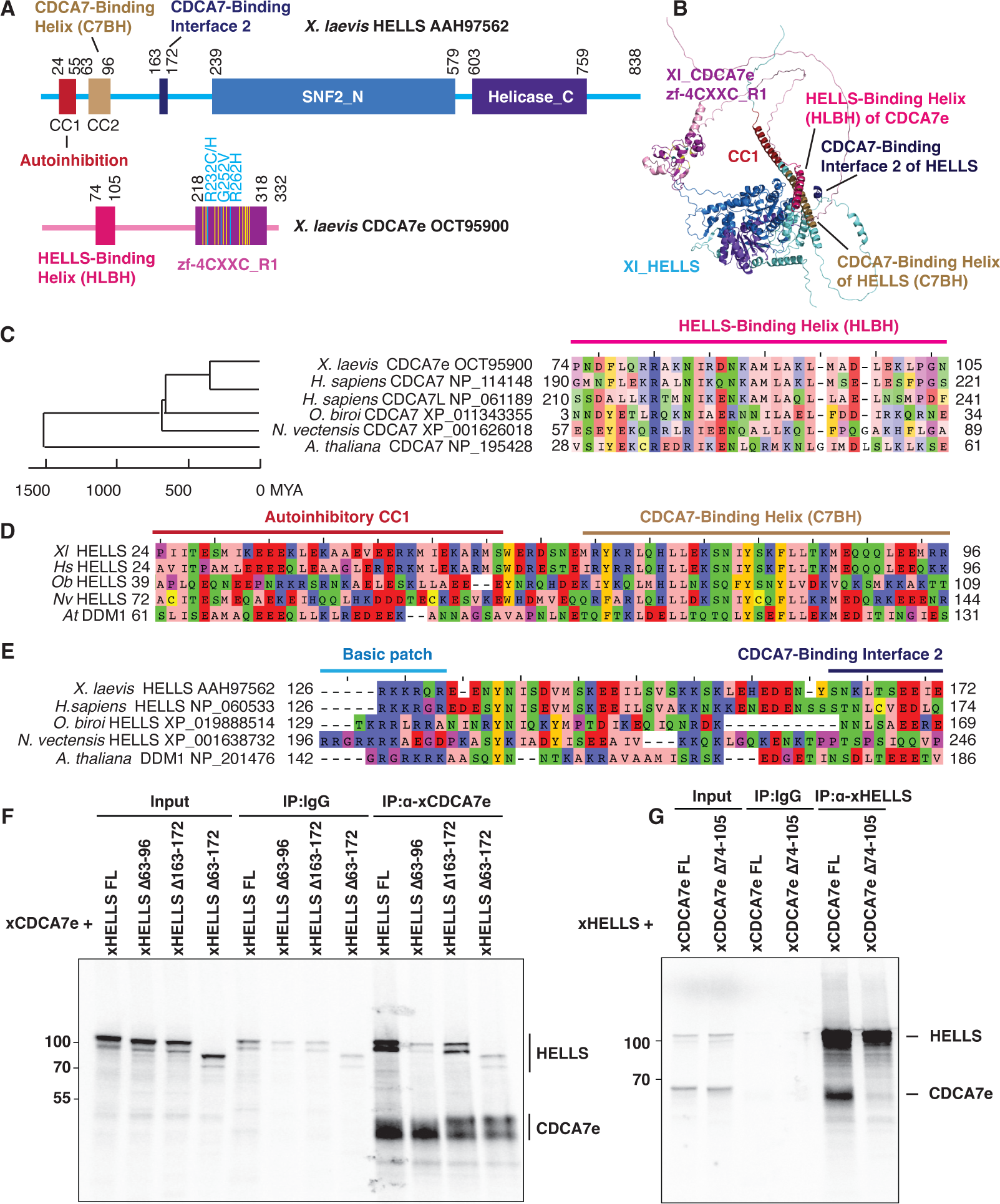
Identification of HELLS-CDCA7 interaction interface. **(A)** Schematics of *X. laevis* HELLS and CDCA7e. Positions of the signature 11 conserved cysteine residues and 3 ICF disease-associated mutations in CDCA7e are marked in yellow and cyan, respectively. CC1 is a coiled-coil domain important for autoinhibition. **(B)** The best predicted structure model of *X. laevis* HELLS-CDCA7e complex by AF2. **(C)** Sequence alignment of the putative HELLS/DDM1-binding interface of CDCA7. **(D)** Sequence alignment of the putative CDCA7-binding interface 1 in HELLS/DDM1. **(E)** Sequence alignment of the putative CDCA7-binding interface 2 in HELLS. **(F)** Immunoprecipitation by control IgG or anti-CDCA7e antibodies from *Xenopus* egg extracts containing ^35^S-labeled wild-type or deletion mutant of *X. laevis* HELLS and CDCA7e. **(G)** Immunoprecipitation by control IgG or anti-HELLS antibody from *Xenopus* egg extracts containing ^35^S-labeled HELLS and wild-type or Δ74-105 deletion mutant of CDCA7e. Autoradiography is shown in F and G.

To experimentally validate these HELLS-CDCA7 binding interfaces, ^35^S-labeled *X. laevis* HELLS or CDCA7e proteins with or without these segments were incubated with *Xenopus* egg extracts to allow for binding to endogenous HELLS/CDCA7e proteins. Co-immunoprecipitation experiments demonstrate that deleting the first predicted CDCA7-binding interface of HELLS (aa 63-96) abolished HELLS-CDCA7e interaction, whereas deleting the second interface of HELLS (aa 163-172) also reduced CDCA7e binding, albeit to a lesser extent (Fig. 4F). This result suggests that the N-terminal CC2 of HELLS acts as a critical CDCA7-binding interface.

Conversely, deleting the predicted HELLS-binding interface in CDCA7e (aa 74-105) abolished HELLS interaction (Fig. 4G). The result was also confirmed by using full-length or truncated versions of recombinant FLAG-tagged CDCA7e (fig. S8); all mutants lacking the N-terminal alpha helix abolished HELLS binding, whereas the N-terminal portion that includes this alpha helix but lacks zf-4CXXC_R1 retains robust HELLS binding. Altogether these data support the AF2 predicted model in which CDCA7 and HELLS interact via their evolutionarily conserved N-terminal helices. We name these helices in CDCA7 and HELLS respectively HLBH (<u>H</u>E<u>L</u>LS-<u>b</u>inding <u>h</u>elix) and C7BH (<u>C</u>DCA<u>7</u>-<u>b</u>inding <u>h</u>elix).

### CDCA7 recruits HELLS to hemimethylated DNA

The experiments above showed that HELLS and CDCA7 are enriched on chromatin with hemimethylated DNA (Fig. 1), and that CDCA7 directly binds to hemimethylated DNA (Fig. 2 and 3). To test if HELLS accumulation onto hemimethylated DNA depends on CDCA7, unmethylated or hemimethylated DNA beads were incubated with mock IgG-depleted (ΔMOCK) or CDCA7e-depleted (ΔCDCA7e) interphase egg extracts. Depletion of CDCA7e did not co-deplete HELLS from egg extracts, but dramatically reduced the binding of HELLS to hemimethylated DNA (Fig. 5A). Furthermore, when ^35^S-labeled HELLS was incubated with egg extracts, it preferentially bound to hemimethylated DNA over unmethylated DNA (Fig. 5B). This hemimethylated DNA-specific binding was abolished by CDCA7 depletion or deleting the CDCA7-binding helix from HELLS (C7BH: Δ63-96) (Fig. 5B, Table S1). Based on these observations, we conclude that CDCA7 recruits HELLS to the hemimethylated DNA.

**Fig. 5.**
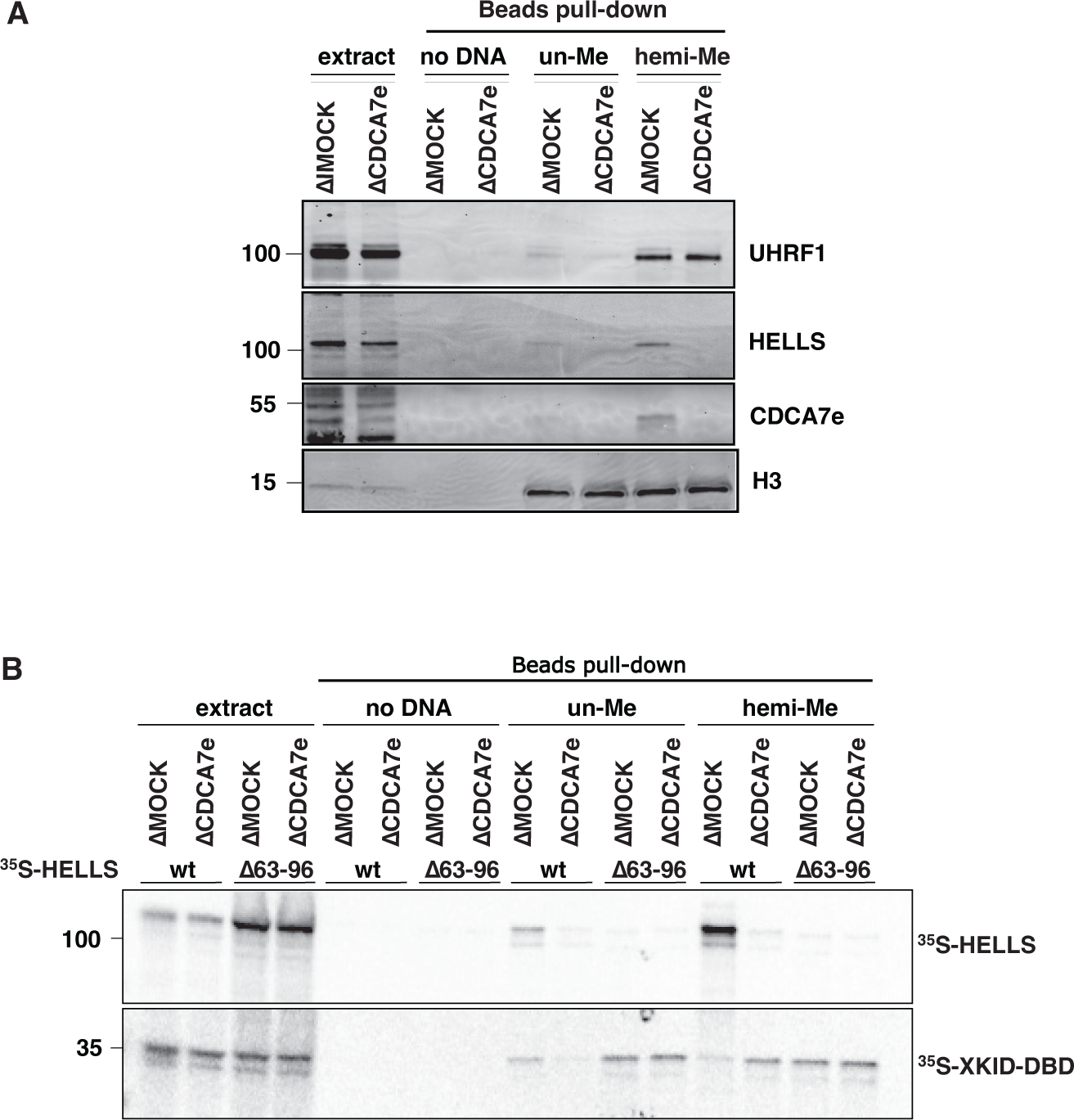
CDCA7 recruits HELLS to hemimethylated DNA. **(A)** Beads coated with unmethylated or hemimethylated DNA (pBluescript) were incubated with interphase *Xenopus* egg control mock IgG-depleted extracts (ΔMOCK) or CDCA7e-depleted extracts (ΔCDCA7e) for 30 min. Beads were isolated and analyzed by western blotting**. (B)** ^35^S-labeled HELLS or HELLS Δ63-96 was incubated with beads coated with 200 bp unmethylated or hemimethylated DNA. Beads were isolated and associated ^35^S-labeled proteins were visualized by autoradiography. Nonspecific DNA-binding protein Xkid DNA-binding domain (Xkid-DBD) was used as a loading control.

### The role of HELLS and CDCA7 in UHRF1-mediated histone H3 ubiquitylation

Studies using ICF patient-derived cells and cell lines, as well as targeted depletion/knockout in culture cells, suggested that HELLS and CDCA7 are especially required for maintaining DNA methylation at heterochromatic, late-replicating regions (*15, 39, 49, 58*). It was also suggested that HELLS/DDM1-dependent methylation is mediated by DNMT1/MET1 (plant DNMT1) (*25, 59*). However, we did not detect any measurable impact of CDCA7e or HELLS depletion on maintenance DNA methylation of sperm or erythrocyte nuclei in *Xenopus* egg extracts as monitored by the incorporation of *S*-[methyl-^3^H]-adenosyl-L-methionine (*35*) (fig. S9). The apparent absence of a role for HELLS and CDCA7e in bulk maintenance DNA methylation could be explained by their function in replication-uncoupled maintenance methylation specifically, which is mediated by UHRF1-dependent H3 ubiquitylation (*37*). Indeed, it has been shown in HeLa cells that HELLS facilitates UHRF1-mediated H3 ubiquitylation (*25*), and promotes the replication-uncoupled maintenance methylation at late-replicating regions (*39*). However, no obvious effect of CDCA7e or HELLS depletion on H3 ubiquitylation was observed when hemimethylated DNA beads were exposed to egg extracts (fig. S10A), even after inducing nucleosome assembly on hemimethylated DNA by preincubating beads in egg extract lacking maintenance methylation (fig. S10B and C). The failure to detect a requirement for CDCA7e and HELLS in H3 ubiquitylation on hemimethylated DNA beads could arise from the specific chromatin environment established on DNA beads in egg extract; nucleosome density as well as the presence or absence of specific histone variants and/or modifications (*60, 61*) are likely specific to the use of an exogenous DNA substrate and conceivably impact the requirement for CDCA7e and HELLS in H3 ubiquitylation.

Therefore, we next attempted to examine the potential role of CDCA7e and HELLS in H3 ubiquitylation on native chromatin after DNA replication. For this purpose, we first induced the accumulation of hemimethylated CpG on sperm nuclei by replicating sperm chromatin in the presence of mDPPA3 (Fig. 6A). These sperm nuclei containing hemimethylated DNA were subsequently transferred to fresh egg extracts with or without aphidicolin, which inhibits DNA replication. As expected, UHRF1 readily and transiently associated with these chromatin substrates and promotes H3 ubiquitylation even in the presence of aphidicolin, demonstrating that UHRF1-mediated H3 ubiquitylation was uncoupled from DNA replication (Fig. 6A). In this experimental context, depletion of CDCA7 or HELLS mildly reduced H3 ubiquitylation and DNMT1 association (Fig. 6B and C). Altogether, these results are in line with the idea that CDCA7 recruits HELLS to hemimethylated chromatin to facilitate replication-uncoupled maintenance methylation.

**Fig. 6.**
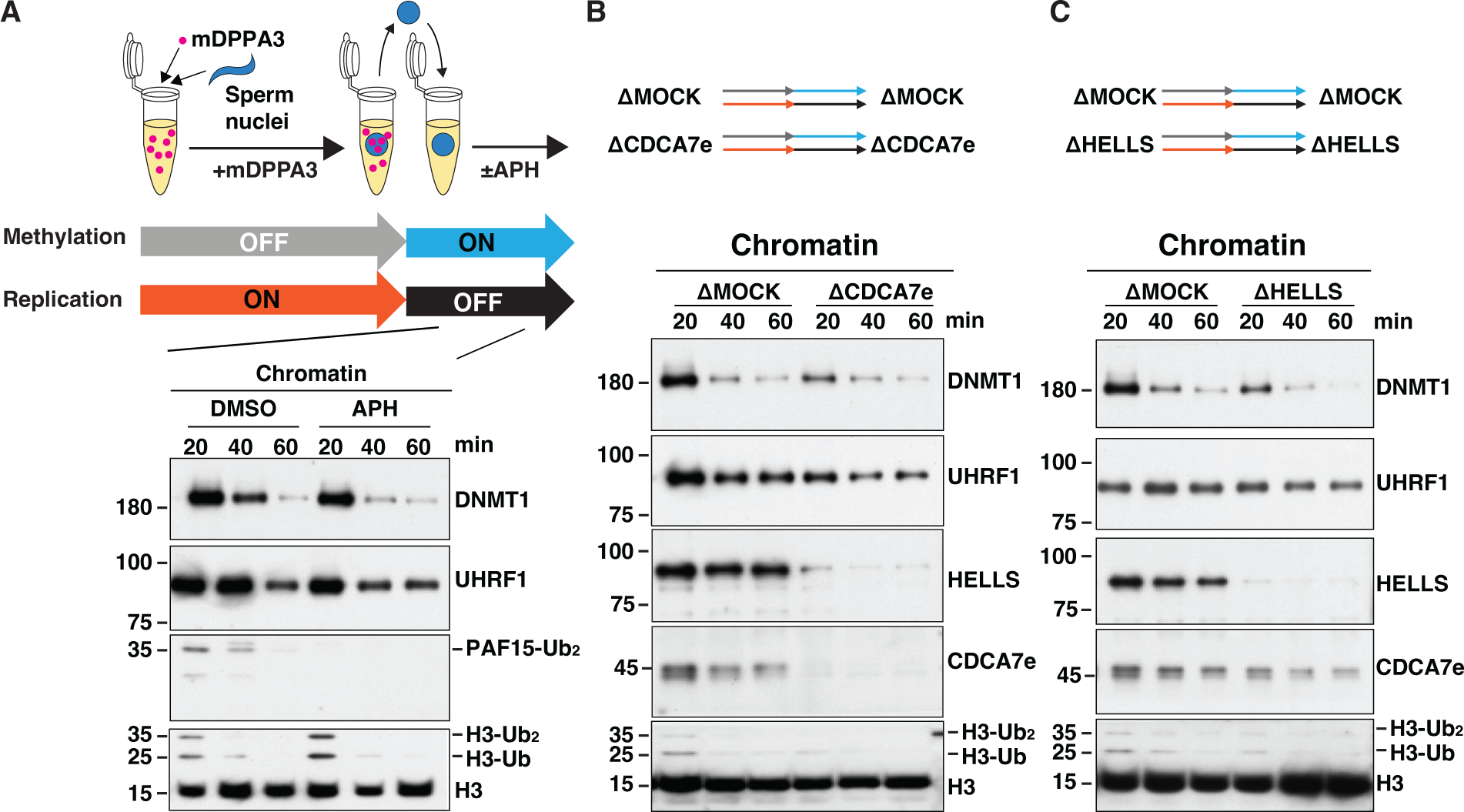
CDCA7e and HELLS regulate replication-uncoupled maintenance DNA methylation. **(A)** *Xenopus* sperm nuclei were incubated for 120 min in interphase *Xenopus* egg extract in the presence of 0.5 µM recombinant mDPPA3. Chromatin was isolated and reincubated in interphase egg extract in the presence or absence of 150 μM aphidicolin (APH). **(B, C).** Sperm nuclei were incubated for 120 min in Mock-depleted extracts or either CDCA7e-depleted **(B)** or HELLS-depleted **(C)** extracts supplemented with mDPPA3. Chromatin was isolated and reincubated in Mock-depleted and either CDCA7e-depleted **(B)** or HELLS-depleted **(C)** extracts in the presence of aphidicolin. Chromatin was then isolated at indicated time points and chromatin-bound proteins were analyzed by western blotting using indicated antibodies.

## Discussion

Among several SNF2-family ATPases that can remodel nucleosomes, HELLS/DDM1 plays a unique role in DNA methylation (*28*). It has also been reported that HELLS promotes replication-uncoupled maintenance DNA methylation by facilitating histone H3 ubiquitylation (*25*). Our present study revealed a previously missing molecular link between HELLS and the maintenance methylation pathway; CDCA7, which recruits HELLS to hemimethylated CpG via its unique zf-4CXXC_R1 domain.

Assisted by AF2 structural prediction, we demonstrated that two evolutionarily conserved alpha helices at the N-terminal regions of CDCA7 and HELLS are responsible for their interaction. It has been shown that HELLS on its own is catalytically inactive (*27, 62*). Deleting the N-terminal alpha helix CC1 of human HELLS preceding the CDCA7-binding helix (C7BH/CC2) activates the ATPase and nucleosome remodeling activities of HELLS (*57*). Similarly, the N-terminal region of *Arabidopsis* DDM1 harboring CC1 and CC2 form an autoinhibitory (AutoN) domain (Fig. 4D) (*48*). Consistent with its proposed autoinhibitory function, the AF2 models predict that the highly acidic CC1 of *Arabidopsis* DDM1 associates with the basic cleft that captures DNA on the nucleosome core particle (NCP) (fig. S6G-I) (*48*); this CC1 placement should interfere with DDM1 binding to the nucleosome. Intriguingly, the AF2 model predicts that the binding of CDCA7 is insufficient to affect CC1 association with the DNA-binding cleft of DDM1 (fig. S6G). It is thus possible that the plant CDCA7 recruits DDM1 to hemimethylated DNA but is not essential for DDM1 activation, although it remains to be tested if the plant CDCA7 binds DDM1. For animal HELLS homologs, CC1 and CC2 are predicted to form a long continuous helix (Fig. 4 and fig. S6C-F), while the acidic feature of the autoinhibitory CC1 is evolutionarily conserved (Fig. 4D). Future studies are needed to test whether binding of CDCA7 activates HELLS/DDM1 by displacing the CC1 from the DNA-binding cleft.

While mutations of DNMT3B (ICF1), ZBTB24 (ICF2), CDCA7 (ICF3) and HELLS (ICF4) cause ICF syndromes, the genomic DNA methylation pattern in the *de novo* DNA methyltransferase-defective ICF1 patient cell lines are distinct from ICF2-4 cell lines, in which CpG-poor regions with heterochromatin features are particularly hypomethylated (*58*). This observation potentially indicates the importance of the ZBTB24-CDCA7-HELLS axis, but not *de novo* DNA methylation, to establish stably inherited DNA methylation at these regions.

Additionally, coevolution analysis demonstrated that CDCA7 and HELLS have stronger evolutionary links to DNMT1 than to DNMT3 (*28*). These findings suggested a function for CDCA7 and HELLS in DNA maintenance methylation at hemimethylated DNA, which are now consolidated by our demonstration that CDCA7 zf-4CXXC_R1 domain specifically recognizes hemimethylated CpG, the substrate of DNMT1. The fact that ICF disease-associated mutations in CDCA7 abolish its hemimethylated DNA binding supports the functional importance of hemimethylation detection by CDCA7.

Since DNA methyltransferases cannot methylate DNA on the NCP (*41–45*), we previously postulated that CDCA7-HELLS promotes DNA methylation by sliding and exposing DNA from the NCP for DNA methyltransferases (*27*). Our biochemical study showed that the zf-4CXXC_R1 domain of CDCA7 can recognize a hemimethylated CpG when placed at the linker DNA but not at a position within the NCP. However, since the positioning of the hemimethylated CpG was not tested exhaustively, we cannot rule out the possibility that the zf-4CXXC_R1 domain binds hemimethylated CpG within the NCP at a different location. Indeed, our cryo-EM structure suggests that the zf-4CXXC_R1 domain recognizes hemimethylated CpG in the major groove of B-form DNA, without requiring any apparent contact with the minor groove. This contrasts with the SRA domain of UHRF1, which extensively engages both the major and minor grooves (*30–32*). As histones contact the minor groove all along the NCP (*63*), it is tempting to speculate that CDCA7 may be more amenable to detecting hemimethylated CpGs in the context of nucleosomes than UHRF1. Although not detected in our structure at the current resolution, it is also possible that specific recognition of hemimethylated CpG by CDCA7 may require DNA distortion in a way that is impossible on the NCP, similar to UHRF1 and DNMT1 (*30-32, 64*). While detailed structural and mechanistic understanding requires further investigation, we envision that the recruitment and activation of HELLS to hemimethylated DNA via CDCA7 unwraps DNA from the NCP, and may additionally increase the accessibility of the histone H3 N-terminal tail that otherwise associates with linker DNA (*65–68*), thereby promoting its recognition by UHRF1 (*69, 70*). In this way, the binding of hemimethylated CpGs by CDCA7 may promote methylation within DNA normally found within the NCP.

We note several limitations in this study. First, since it has been shown that binding of UHRF1 to histone H3 di- or tri-methylated at lysine 9 (H3K9me2/3) facilitates DNA methylation at heterochromatin (*39, 71*), the role of CDCA7 in maintenance methylation in the context of heterochromatic nucleosomes remains to be tested. Second, although this study focused on the role of CDCA7 in maintenance methylation, it is possible that hemimethylated DNA sensing by CDCA7 also plays an important role outside of this process. Indeed, DNA methylation in insects is largely associated with gene bodies and not with heterochromatic transposable elements (*72–75*), apparently contradicting the suggested specialized role of CDCA7-HELLS in maintaining DNA methylation at heterochromatin. It will be important to test the functional significance of CDCA7-hemimethylated CpG binding in other processes where HELLS and/or CDCA7 play roles, such as DNA repair, resolution of DNA-RNA hybrids, and macroH2A deposition (*49, 62, 76-82*). Third, the low resolution of our current cryo-EM structure of the CDCA7-nucleosome complex prevented us from dissecting the structural basis for hemimethylated CpG recognition by CDCA7 at atomic resolution. Fourth, although our data clearly show that CDCA7 selectively binds to DNA with a single hemimethylated CpG over unmethylated or symmetrically methylated CpG, further investigations are needed to test if CDCA7 has more optimized substrates. The binding may be affected by DNA sequence, density and spacing of hemimethylated CpG, or other modifications, such as 5-hydroxymethylcytosine.

## Materials and Methods

### *Xenopus* egg extracts

At the Rockefeller University, *Xenopus laevis* was purchased from Nasco (female, LM00535MX) or Xenopus 1 (female, 4270; male, 4235); and all vertebrate animal protocols (20031 and 23020) followed were approved by the Rockefeller University Institutional Animal Care and Use Committee. In Fig. 1A; Fig. 2A; Fig. 4F and G; Fig. 5; fig. S9 and fig. S10, freshly prepared crude cytostatic factor (CSF) metaphase-arrested egg extracts were prepared as previously published (Murray, 1991). To prepare interphase extracts, 0.3 mM CaCl_2_ was added to CSF extract containing 250 ng/μl cycloheximide.

At the Institute of Medical Science, University of Tokyo, *X. laevis* was purchased from Kato-S Kagaku and handled according to the animal care regulations at the University of Tokyo. In Fig. 1B, Fig. 6, fig. S2 and fig. S7, clarified cytoplasmic extracts were used. Crude interphase egg extracts were prepared as described previously (*37, 83*), supplemented with 50 μg/ml cycloheximide, 20 μg/ml cytochalasin B, 1 mM dithiothreitol (DTT), 2 μg/ml aprotinin, and 5 μg/ml leupeptin and clarified by ultracentrifugation (Hitachi, CP100NX, P55ST2 swinging rotor) for 20 min at 48,400× g. The cytoplasmic extracts were aliquoted, frozen in liquid nitrogen, and stored at –80°C. The clarified cytoplasmic extracts were supplemented with an energy regeneration system (2 mM ATP, 20 mM phosphocreatine, and 5 μg/ml creatine phosphokinase).

### Chromatin isolation

*Xenopus* sperm nuclei (3000-4000 per μl) was added to interphase extract and incubated at 22 °C. Extract was diluted five- to ten-fold in chromatin purification buffer (CPB; 50 mM KCl, 5 mM MgCl_2_, 2 % sucrose, 20 mM HEPES-KOH, pH 7.6) supplemented with 0.1 % Nonidet P-40 (NP-40). With the exception of Fig. 1A, CPB was additionally supplemented with 2 mM NEM and 0.1 mM PR-619. Diluted extracts were layered onto a CPB-30% sucrose cushion and centrifuged at 15,000× *g* for 10 min at 4 °C. The chromatin pellet was recovered in 1x Laemmli sample buffer, boiled and Western blotting was performed against the indicated proteins.

### Antibodies and western blotting

*Xenopus* CDCA7e, HELLS, PAF15, DNMT1 and UHRF1 were detected with rabbit polyclonal antibodies previously described (*27, 35, 37*). Rabbit polyclonal histone H3 antibody (ab1791) was purchased from Abcam. Rabbit polyclonal histone H4 antibody (Cat #61521) was purchased from Active Motif. In Fig. 1A; Fig. 2A; Fig. 5; and fig. S10, antibodies were used in LI-COR Odyssey blocking buffer at the following dilutions: affinity purified anti-CDCA7e (2 µg/ml), affinity purified anti-HELLS (3.5 µg/ml), anti-UHRF1 serum (1:500), anti-H3 (1:1000); anti-H4 (1:1000). Primary antibodies were detected with IRDye^®^ secondary antibodies (Cat #926-32211; Cat #926-68070, LI-COR BioSciences) and subsequently imaged and quantified on an Odyssey Infrared Imaging System. In Fig. 1B, Fig. 6, fig. S2 and fig. S7, anti-DNMT1 (1:500) and anti-UHRF1 (1:500) sera were used in 5% Milk in PBS-T; anti-PAF15 (1:500), anti-CDCA7e (1:500) and anti-HELLS (1:500) sera were used in Sol 1 (Toboyo, Can Get Signal® Immunoreaction Enhancer Solution). Primary antibodies were detected with HRP-conjugated secondary antibodies (rabbit IgG-, Protein A-, or mouse IgG-conjugated with HRP, Thermo Fisher Scientific) and ECL detection reagent (Amersham). After exposure to the wrapped membrane, X-ray film was developed.

### Immunodepletion

To immunodeplete CDCA7e or HELLS from extracts used for DNA beads pull-down experiments, 37.5 µg affinity purified anti-CDCA7 or anti-HELLS antibodies was coupled to 150 µl Protein A Dynabeads (Thermo Fisher Scientific) and used to deplete 100 µl extract at 4 °C for 45 min. To immunodeplete DNMT1 from extract used for chromatin isolation experiments, 85 µl serum was coupled to 25 µl Protein A Dynabeads (Thermo Fisher Scientific) and used to deplete 33 μL extract in three separate rounds at 4 °C, each for 1 h. Prior to depletion, antibody-coupled Dynabeads were washed extensively in sperm dilution buffer (5 mM HEPES, 100 mM KCl, 150 mM sucrose, 1mM MgCl_2_, pH 8.0). To immunodeplete CDCA7e or HELLS from extract used for chromatin isolation experiments, 170 µl of antiserum was coupled to 40 µl of recombinant Protein A Sepharose (rPAS, GE Healthcare). Antibodies bound beads were washed extensively in CPB and supplemented with 4 µl fresh rPAS. Beads were split into two portions, and 100 µl of extract was depleted in two rounds at 4°C, each for 1 h. Mock depletion was performed using purified preimmune rabbit IgG (Sigma-Aldrich).

### Immunoprecipitations

For coimmunoprecipitation from *Xenopus* egg extracts, anti-HELLS and anti-CDCA7e antibodies (25 μg) were coupled to 100 μl Protein A Dynabeads for 1 h at RT. Antibodies were crosslinked to the beads with Pierce^TM^ BS_3_ (Thermo Fisher Scientific), following the manufacturer’s protocol. Antibody beads were washed extensively in sperm dilution buffer (5 mM HEPES, 100 mM KCl, 150 mM sucrose, 1 mM MgCl_2_, pH 8.0). To test CDCA7 and HELLS interactions, CDCA7e or HELLS wildtype or interface mutants were expressed and radiolabeled with EasyTag^TM^ L-[^35^S]-Methionine (Perkin Elmer) using the TnT Coupled Reticulocyte Lysate System (Promega) according to the manufacturer’s instructions. HELLS and CDCA7 mutants were cloned into pCS2 vector by Gibson assembly. Immunoprecipitation was performed in 50 μl interphase egg extracts supplemented with 250 ng/ μl cycloheximide and 0.08 μl of the indicated ^35^S-labeled CDCA7 and HELLS TnT lysates per μl of extract. Extract was added to the beads and incubated on ice for 1 h with flicking every 20 min. The extract was diluted with 10 volumes CSF-XB (100 mM KCl, 1 mM MgCl_2_, 50 mM sucrose, 5 mM EGTA, and 10 mM HEPES, pH 8.0) and beads were recovered on a magnet. Beads were washed and recovered three times with 150 μl CSF-XB with 0.1 % Triton X-100. Beads were resuspended in 1x Laemmli buffer, boiled and supernatants were resolved by SDS-PAGE. Gels were fixed in fixative (1:2:7 glacial acetic acid:methanol:H_2_O), dried and exposed on a PhosphorImager screen. Control immunoprecipitation was performed using purified preimmune rabbit IgG (Sigma-Aldrich).

### DNA pull-down assays

To generate hemimethylated pBlueScript DNA substrates, a PCR-linearized pBlueScript template was methylated by the CpG methyltransferase M.SssI according to manufacturer’s protocol (Cat #EM0821, Thermo Fisher Scientific). DNA synthesis across the methylated linearized pBlueScript template was subsequently performed in Q5^®^ High-Fidelity 2X Master Mix (New England Biolabs, Inc.) using a 5’ biotinylated primer (5’-/5Biosg/CGTTCTTCGGGGCGAAAACTCTCAAGG −3’) purchased from Integrated DNA Technologies. The reaction mix was purified using the QIAquick PCR purification kit (QIAGEN) and the resultant hemimethylated DNA product was subsequently purified from the reaction mix by conjugation to streptavidin M280 dynabeads (Invitrogen). DNA was coupled to streptavidin beads at ∼2 μg DNA/5 μl bead slurry in bead coupling buffer (50 mM Tris-Cl, 0.25 mM EDTA, 0.05% Triton X-100, pH 8.0) supplemented with 2.5% polyvinyl alcohol and 1.5 M NaCl for at least 2 h at RT. After conjugation, DNA-streptavidin beads were collected and incubated in 50 mM Tris-Cl, 0.25 mM EDTA, 0.05% Triton X-100 with 1 mM biotin for at least 30 min. DNA beads were extensive washed in sperm dilution buffer (5 mM HEPES, 100 mM KCl, 150 mM sucrose, 1 mM MgCl_2_, pH 8.0) prior to performing any pull-down assay. For nonmethylated BlueScript DNA substrates, the above protocol was performed using unmethylated linearized pBlueScript template during DNA synthesis. Fully-methylated pBlueScript DNA substrates were generated by methylating the nonmethylated pBlueScript DNA substrates by CpG methyltransferase M.SssI (Thermo Fisher Scientific) prior to DNA-bead conjugation. 200 bp ultramers with Widom 601 nucleosome positioning sequence (Table S1) (*84*) were purchased from Integrated DNA Technologies and conjugated to streptavidin M280 Dynabeads as described above at ∼1 μg DNA/5 μl bead slurry. Methylation status of all DNA substrates was confirmed by restriction digest with BstUI (New England Biolabs, Inc.).

For DNA bead pull-downs analyzed by western blot (Fig. 2A, Fig. 5A, fig. S10) DNA beads were incubated in interphase *Xenopus* egg extract. The extract was diluted with 10 volumes CSF-XB and recovered on a magnet for 5 min at 4 °C. Beads were washed and recovered three times with 150 μl CSF-XB with 0.1% Triton X-100. Beads were resuspended in 1x Laemmli buffer, boiled and supernatants were resolved by SDS-PAGE. Western blotting was performed against the indicated proteins. To assess protein binding by autoradiography (Fig. 2B; Fig. 4F-G and Fig. 5B), indicated proteins were expressed and radiolabeled with EasyTag^TM^ L-[^35^S]-Methionine (Perkin Elmer) using the TnT Coupled Reticulocyte Lysate System (Promega) according to the manufacturer’s instructions. ^35^S-labeled Xkid (*85*) or Xkid-DNA binding domain was used as a DNA loading control. The C-terminally GFP tagged DNA-binding domain of Xkid (Xkid-DBD, amino acids 544–651) was cloned into pCS2 vector by Gibson assembly. To assess the recruitment of HELLS to hemimethylated DNA in egg extract using autoradiography (Fig. 5B), DNA bead pull-down was performed in Interphase *Xenopus* egg extract supplemented with 0.1 μl ^35^S-labeled HELLS and 0.03 μl ^35^S-labeled Xkid-DBD per μl of extract. Beads were washed and recovered three times with 150 μl CSF-XB with 0.1 % Triton X-100. Beads were resuspended in 1x Laemmli buffer, boiled and supernatants were resolved by SDS-PAGE. Gel was fixed in fixative (1:2:7 glacial acetic acid:methanol:H2O), dried and exposed on a PhosphorImager screen. To assess the in vitro binding of CDCA7e ICF mutants to hemimethylated DNA by autoradiography (Fig. 2B), DNA bead pull-down was performed in binding buffer (10 mM HEPES, 100 mM NaCl, 0.025 % Triton X-100, 0.25 mM TCEP, pH 7.8) supplemented with 0.2 μl ^35^S-labeled CDCA7 and 0.05 μl ^35^S-labeled Xkid per μl binding buffer. Beads were washed and recovered three times with binding buffer supplemented with 0.1% Triton X-100. Beads were resuspended in 1x Laemmli buffer, boiled and resolved by gel electrophoresis. All DNA pull-downs were performed at 20 °C.

### Detection of DNA methylation maintenance in *Xenopus* egg extract

DNA methylation of replicating sperm or erythrocyte nuclei in egg extract was assayed by the incorporation of ^3^H-SAM (S-[methyl-^3^H]-adenosyl-L-methionine; Perkin Elmer, NET155H). Demembranated sperm nuclei were prepared as published previously (*86*). Erythrocyte nuclei were prepared from blood collected from dead adult male *Xenopus laevis* frogs that were sacrificed for testis dissection, following the protocol published previously (*87*), with the addition of an extra dounce homogenization step prior to pelleting the nuclei over the 1M sucrose cushion. Erythrocyte nuclei were stored at −20 ° C in 50% glycerol STMN buffer (10 mM NaCl, 10 mM Tris pH 7.4, 3 mM MgCl_2_, 0.5% NP-40). Sperm or erythrocyte nuclei were replicated in cycling egg extract (3000 nuclei/μl extract) supplemented with 250 ng/μl cycloheximide and 0.335 μM ^3^H-SAM (82.3 Ci/mmol) for 1 h at 20 °C. Replication was inhibited by the addition of 200 nM of recombinant GST-tagged nondegradable geminin (fig. S9, an expression plasmid provided by W. Matthew Michael) (*54*) or 500 nM of 6His-geminin (fig. S2, a gift from Tatsuro Takahashi). The reaction was stopped by the addition of 9 volumes of CPB. Genomic DNA was purified using a Wizard Genomic DNA Purification Kit (Promega) according to the manufacturer’s instructions. Chromatin pellets were resuspended in scintillation fluid (ScintiVerse; Thermo Fisher Scientific) and quantified using a liquid scintillation counter (Perkin Elmer, Tri-Carb^®^ 2910 TR).

### Protein purification

For FLAG×3-tagged full-length mDPPA3 or xCDCA7e expression in insect cells, Baculoviruses were produced using a BestBac v-cath/chiA Deleted Baculovirus Cotransfection kit (Expression system) following the manufacturer’s instructions. Proteins were expressed in Sf9 insect cells by infection with viruses expressing 3xFLAG-tagged mDPPA3 or xCDCA7e for 72 h at 27 °C. Sf9 cells from a 750 ml culture were collected and lysed by resuspending them in 30 ml lysis buffer, followed by incubation on ice for 10 min. A soluble fraction was obtained after centrifugation of the lysate at 15,000× *g* for 15 min at 4 °C. The soluble fraction was incubated for 4 h at 4 °C with 250 µl of anti-FLAG M2 affinity resin equilibrated with lysis buffer. The beads were collected and washed with 10 ml wash buffer and then with 5 ml of EB [20 mM HEPES-KOH (pH 7.5), 100 mM KCl, 5 mM MgCl_2_] containing 1 mM DTT. Each recombinant protein was eluted twice in 250 µl of EB containing 1 mM DTT and 250 µg/ml 3×FLAG peptide (Sigma-Aldrich). Eluates were pooled and concentrated using a Vivaspin 500 (GE Healthcare).

cDNA of human CDCA7 encoding residues 264-371, 235-340 and 264-340 were sub-cloned into modified pGEX4T-3 plasmid (Cytiva) engineered for N-terminal GST and a small ubiquitin-like modifier-1 (SUMO-1) fusion tag (*88*). The protein was expressed *E.coli* strain Rosetta 2 (DE3) (Novagen). The cells were grown at 37 °C in Luria-Bertani medium (LB) containing 50 µg/ml ampicillin and 34 µg/ml chloramphenicol until reaching on optical density of 0.7 at 660 nm, and then cultured in 0.2 mM IPTG for 15 h at 15 °C. The cells were lysed by sonication in 40 mM Tris-HCl (pH 8.0) buffer containing 300 mM NaCl, 0.1 mM DTT (or 0.5 mM TCEP for CDCA7 residues 235-340 and 264-340), 30 µM zinc acetate, 10% (W/V) glycerol and a protease inhibitor cocktail (Nacalai). After removing the debris by centrifugation, the supernatant was loaded onto Glutathione Sepharose 4B (Cytiva). After GST-SUMO tag was removed by SUMO-specific protease, the sample was loaded onto HiTrap Heparin column (Cytiva). Finally, the protein was further purified using size-exclusion chromatography Hiload 26/600 S75 (Cytiva).

### Electrophoresis mobility shift assay

10 µl of samples were incubated for 30 min at 4 °C in a binding buffer [20 mM Tris-HCl (pH 7.5) containing 150 mM NaCl, 1 mM DTT, 0.05 % NP-40 and 10% (w/v) glycerol] and electrophoresis was performed using a 0.5 × Tris-Acetate buffer [20 mM Tris-Acetic acid containing 0.5 mM EDTA at constant current of 8 mA for 100 min in a cold room on a 7.5% polyacrylamide gel purchased from Wako (SuperSep^TM^). 0.5, 1.0 and 2.0 equimolar excess of the CDCA7 264-371 were added to the sample solution including 0.5 µM hemi-, full- and un-methylated DNA (upper: 5’-CAGGCAATC<u>X</u>GGTAGATC, lower: 5’-GATCTAC<u>X</u>GGATTGCCTG, where X indicates cytosine or 5-methylcytosine). 3.0, 5.0 and 10.0 equimolar excess of the CDCA7 264-340 and 235-340 were added to the sample solution including the 0.5 µM DNAs.

For analyzing the interaction with reconstituted nucleosomes, 0.5, 1.0, 2.0 and 5.0 equimolar excess of the CDCA7 264-371 or 0.77, 1.54 and 3.85 equimolar excess of Flag×3-xCDCA7WT or Flag×3-xCDCA7R232H were added to 0.1 µM nucleosomes in 10 µl reaction solution (binding buffer: 20 mM Tris-HCl (pH 7.5), 50 mM NaCl, 1 mM DTT, 10 % Glycerol, 0.05 % NP-40) and electrophoresis was performed using a 0.5 × TBE buffer [(45 mM Tris-borate and 1 mM EDTA) at constant current of 10 mA for 95 min in a cold room on a 7.5% polyacrylamide gel. To analyze the interactions, DNA was detected and analyzed by staining with GelRed^TM^ (Wako) and the ChemiDoc XRS system (BIORAD), respectively.

### Nucleosome reconstruction

Recombinant human histone H2A, H2B, H3.1 and H4 proteins were produced in *Escherichia coli* and purified using gel filtration chromatography and cation exchange chromatography as reported previously (*89*). The histone proteins were refolded into a histone octamer. All DNA including a single hemimethylated CpG were based on the Widom601 nucleosome positioning sequence (*84*). For preparation of DNA with a hemimethylated CpG at the 5’-linker, the Widom601 sequence was amplified using the primers (Table S2). For preparation of DNA with a hemimethylated site in the 3’-linker and nucleosomal DNA, the Widom601 sequence was amplified with BsmBI site at the 3’-region and digested by BsmBI (Table S2). The fragment was ligated with oligonucleotides including a single hemimethylated CpG (Table S2). The DNAs were purified with anion-exchange chromatography, HiTrap Q HP (Cytiva). The histone octamers were reconstituted into nucleosome with purified DNAs by salt dialysis method and the nucleosomes were purified with anion-exchange chromatography, HiTrap Q HP. The purified nucleosomes were dialyzed against 20 mM Tris–HCl buffer (pH 7.5), containing 1 mM DTT and 5% glycerol. The nucleosomes were frozen in liquid nitrogen and stored at −80°C.

### Cryo-EM data collection and data processing

3 µL of the human CDCA7_264-371_ in complex with the nucleosome harboring a single hemimethylated CpG in the 3’-linker DNA was applied onto the glow-discharged holey carbon grids (Quantifoil Cu R1.2/1.3, 300 mesh). The grids were plunge-frozen in liquid ethane using a Vitrobot Mark IV (Thermo Fisher Scientific). Parameters for plunge-freezing were set as follows: blotting time, 3 sec; waiting time, 3 sec; blotting force, −10; humidity, 100 %; and chamber temperature, 4 °C. Data was collected at RIKEN BDR on a 300-kV Krios G4 (Thermo Fisher Scientific) with a K3 direct electron detector (Gatan) with BioQuantum energy filter. A total of 4,000 movies were recorded at a nominal magnification of ×105,000 with a pixel size of 0.83 Å, in a total exposure of 60.725 e^−^/Å^2^ per 48 frames with an exposure time of 2.2 sec. The data were automatically acquired by the image shift method of the EPU software (Thermo Fisher Scientific), with a defocus range of −0.8 to −1.6 μm.

### Data processing

All Data were processed using cryoSPARC v4.2.1 and v4.4.0 (*90*). The movie stacks were motion corrected by Patch Motion Correction. The defocus values were estimated from the Contrast transfer function (CTF) by Patch CTF Estimation. Micrographs under 8 Å CTF resolution were cut off by Curate Exposures, and 3,973 micrographs were selected. A total of 1,881,583 particles were automatically picked using Blob Picker. Particles (1,583,471) were extracted using binning state (3.31 Å/pixel) and these particles were subjected to 2D Classifications. Particles were further curated by Heterogeneous Refinement using the maps derived from cryoSPARC *Ab-Initio* Reconstruction as the template. The selected suitable class containing 854,921 particles were classified by several round of Heterogeneous Refinement. Finally, 3D reconstruction using 672,791 particles was performed by Non-uniform-refinement, and a 2.9 Å resolution map was obtained according to the gold-standard Fourier shell correlation (FSC) = 0.143 criterion. For further classification, 3D Variability and 3D Classification were conducted, obtaining the maps at 3.3 Å resolution which includes the map corresponding to the linker DNA region. Non-uniform-refinement was performed using the particles selected by 3D Variability and 3D Classification (82,876 and 94,542 particles, respectively) (*88*) and a 3.2 Å resolution map was obtained (fig. S4).

The focused mask corresponding to linker DNA containing the hemimethylated CpG bound by hCDCA7 was created for Local refinement. Local refinement improved the map at 4.83 Å resolution (154,998 particles) (fig. S5).

The structure of nucleosome moiety was created using PDB ID: 3LZ0. Linker DNA was generated using program Coot (*91*) and the structure of the nucleosome combined with linker DNA was refined with program PHENIX (*92*). A structure model of human CDCA7_264-371_, was generated from AlphFold2 (AF-Q9BWT1-F1). The model was manually fitted to the focused map, taking into account the surface potential of the protein and characteristic C-terminal α-helix of hCDCA7. Details of the data processing are shown in fig. S4, S5 and table S3. The protein structures were visualized using Pymol (The PyMOL Molecular Graphics System, Version 2.2, Schrödinger, LLC.) and UCSF ChimeraX (Version 1.5)

### Prediction of the interactions between HELLS and CDCA7

We collected protein sequences of HELLS and CDCA7 from five species, including *H. sapiens*, *X. laevis*, *O. biroi*, *N. vectensis,* and *A. thaliana*. We then ran AlphaFold2 (version 2.2.2) (*93*) to predict the interactions between HELLS and CDCA7 in the five different species. For each prediction, we selected the best model for further structural analysis. We implemented a cutoff distance of 5 Å between non-hydrogen atoms to extract the interface residues between HELLS and CDCA7. The same cutoff was also applied to compute the pDockQ (*94*) metric for each of the five predictions. A pDockQ of greater than 0.23 indicates an acceptable predicted model, while a pDockQ of greater than 0.5 indicates a confident predicted model (*94*). To evaluate the convergence of the interface, we used MAFFT (*95*) to align the HELLS and CDCA7 protein sequences, respectively. The sequence alignments were visualized using MVIEW (*96*). The protein structures were visualized using Pymol (The PyMOL Molecular Graphics System, Version 2.1, Schrödinger, LLC.)

### Statistical Analysis

Cryo-EM data collection statistics are available in Table S3.

## Acknowledgments

We thank Hideaki Konishi for construction of the plasmid expressing Xkid-DBD, W. Matthew Michael and Tatsuro Takahashi for geminin, Tomohiro Nishizawa and Yongchan Lee for supporting cryo-EM analysis, Jason Banfelder, Bala Jayaraman, and The Rockefeller University High Performance Computing (HPC) Center for their support in computation, members of the Funabiki lab for discussion, Yasuhiro Arimura for comments on the manuscript, and The Rockefeller University Comparative Bioscience Centre for animal husbandry. The cryo-EM experiments were performed at the cryo-EM facility of the RIKEN Center for Biosystems Dynamics Research Yokohama.

## Funding

National Institutes of Health grant R35GM132111 (HF)

National Institutes of Health grant R35GM133780 (LZ)

The Robertson Foundation (LZ)

MEXT/JSPS KAKENHI grant JP19H05740 (MN)

MEXT/JSPS KAKENHI grant JP19H03143 (AN)

MEXT/JSPS KAKENHI grant JP19H05285 (AN)

MEXT/JSPS KAKENHI JP19H05741 (KA)

The Rockefeller University Women & Science Postdoctoral Fellowship (IEW)

## Author contributions

Conceptualization: IW, AN, HF

Methodology: IW, AN, KA, JP, HF

Investigation: IW, AN, MH, QJ, RS, AK, KS, XH, YC, CJ, KA, HF

Visualization: AK, JP, HF

Supervision: AN, MN, LZ, KA, HF

Writing—original draft: HF, IW, AN, KA, JP

Writing—review & editing: HF, IW, AN, KA, QJ

## Data and materials availability

Cryo-EM maps (EMD-38198 and EMD-38199) were deposited to EMDB and will be released upon publication. All data and materials used in the analyses will be available to any researcher for the purposes of reproducing or extending the analyses. All data are available in the main text or the supplementary materials.

---

**Fig. S1.**
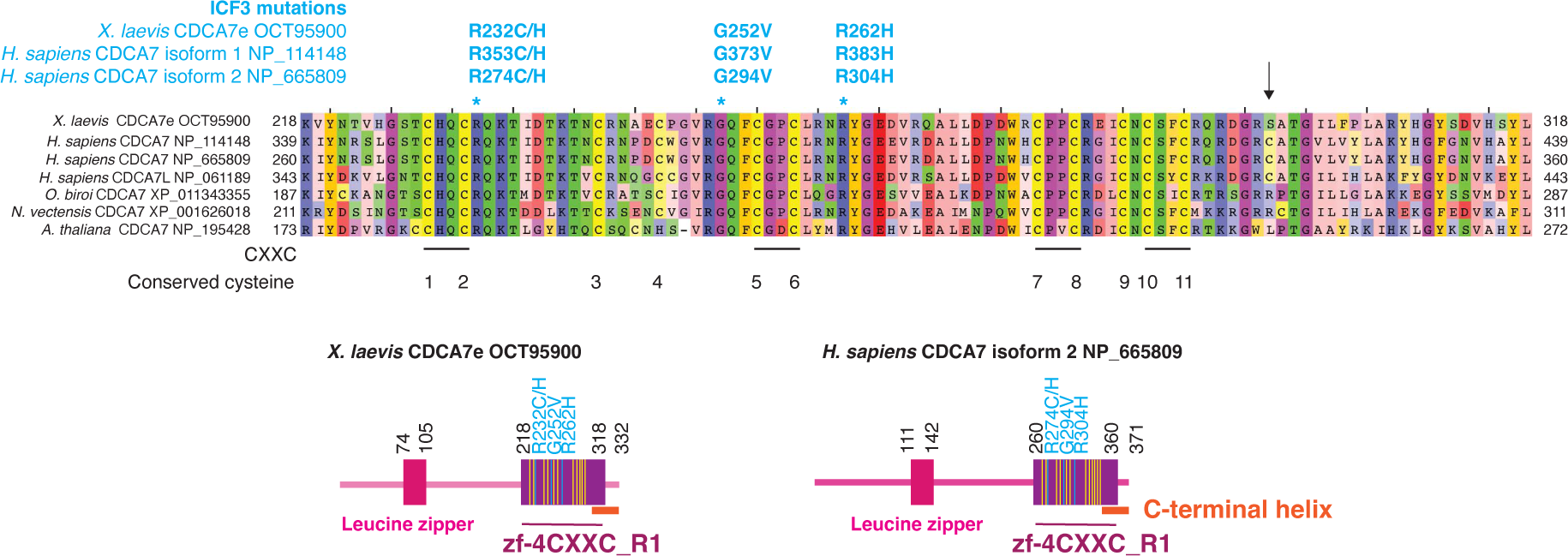
Evolutionary conservation of the zf-4CXXC_R1 domain of CDCA7 homologs. ClustalW multi-sequence alignment of CDCA7 zf-4CXXC_R1 domain, characterized by eleven conserved cysteine (yellow) and three ICF3 patient-associated (cyan) residues. *X. laevis*, *Xenopus laevis* (African clawed frog); *H. sapiens*, *Homo sapiens* (human); *O. biroi*, *Ooceraea biroi* (clonal raider ant); *N. vectensis*, *Nematostella vectensis* (starlet sea anemone); *A. thaliana*, *Arabidopsis thaliana* (thale cress). Amino acid positions of ICF3 associated mutations in *X. laevis* CDCA7e, *H. sapiens* CDCA7 isoform 1 (NP_114148) and the shorter isoform 2 (NP_665809) are indicated. An arrow indicates the position of cysteine 339 of *H. sapiens* CDCA7 isoform 2, the site that was mutated to serine in Fig. 3, fig. S3D, fig. S4 and fig. S5. Schematics showing the domain composition of *X. laevis* CDCA7e and *H. sapiens* CDCA7 isoform 2 are also shown.

**Fig. S2.**
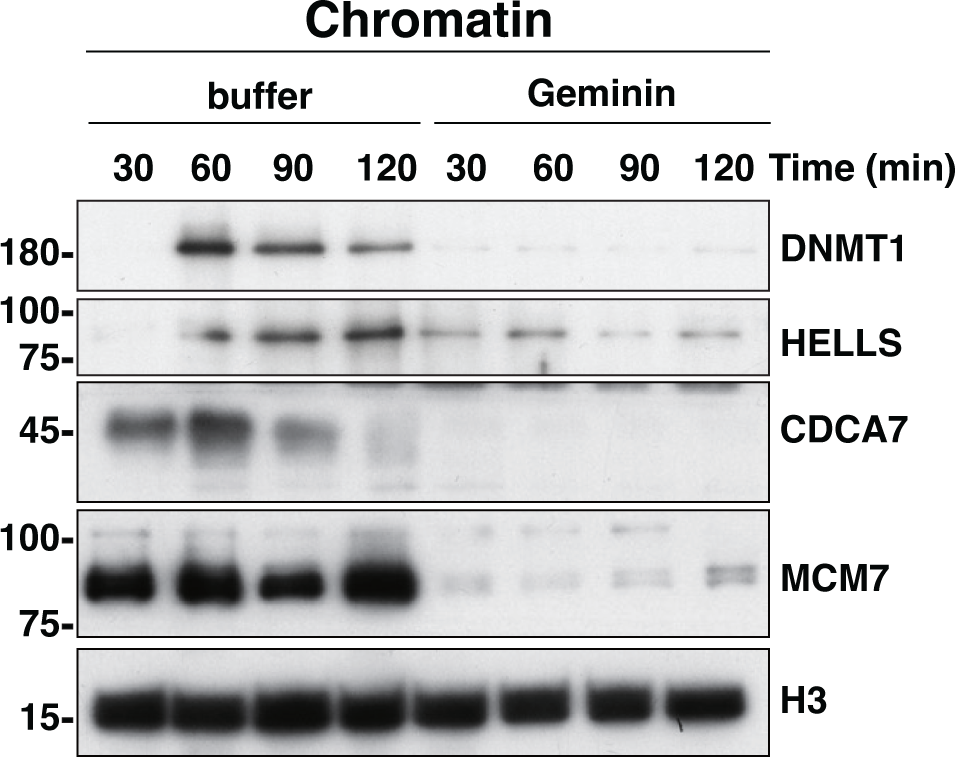
DNA replication promotes chromatin association of CDCA7e and HELLS. **(A)** *X. laevis* sperm nuclei were incubated with interphase *Xenopus* egg extracts in the presence or absence of 0.5 µM recombinant geminin. At each indicated time point, chromatin was isolated and analyzed by western blotting.

**Fig. S3.**
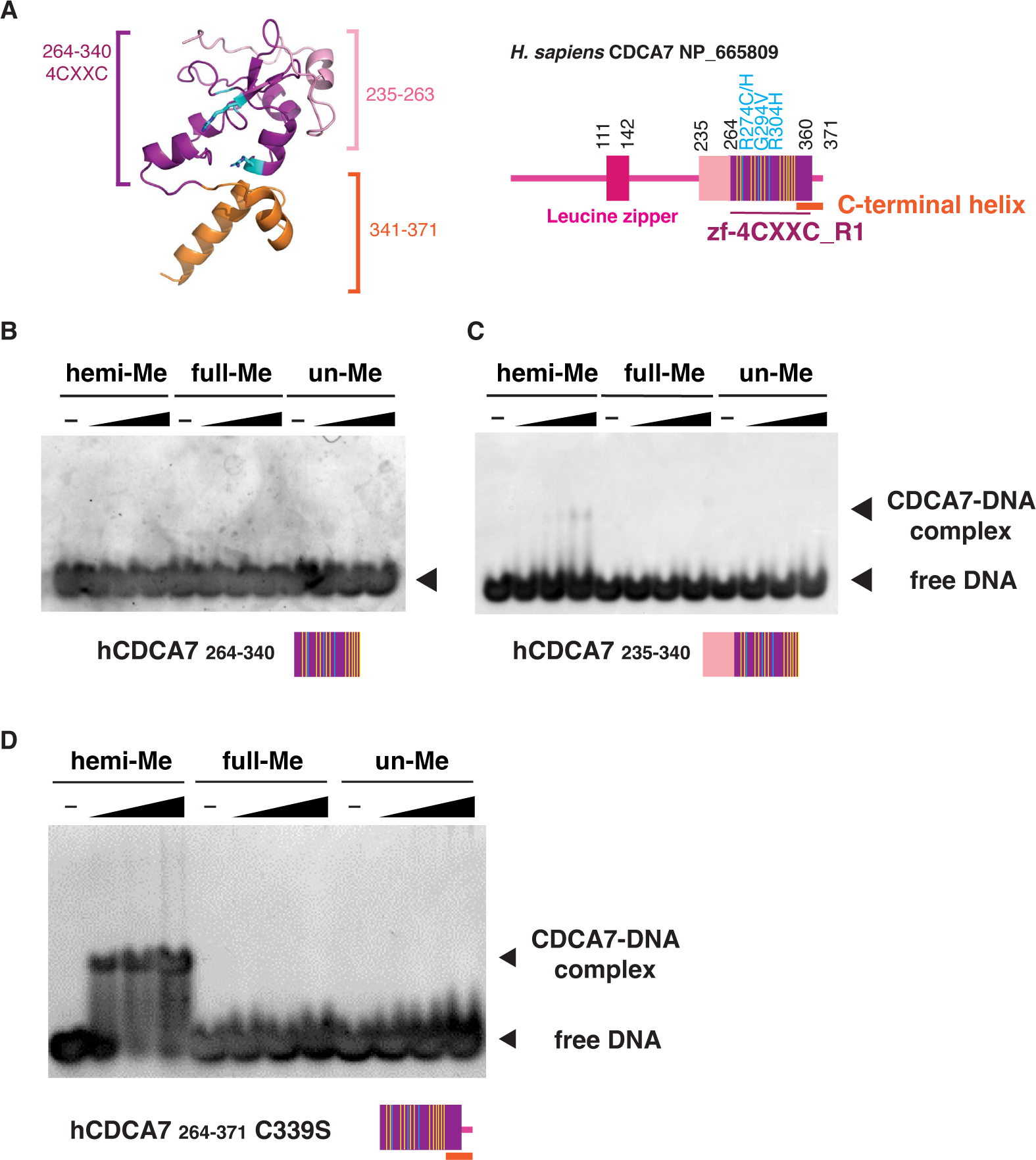
Characterization of the minimum hemimethylated DNA-binding domain of human CDCA7. **(A)** AlphaFold2-modeled structure of *H. sapiens* zf-4CXXC_R1 domain and schematic of full-length *H. sapiens* CDCA7. Yellow lines indicate the position of conserved cysteine residues. Orange bar indicates the conserved C-terminal helix. **(B-D)** Native gel electrophoresis mobility shift assay for detecting the interaction of hCDCA7 _264-340_ **(B)** hCDCA7 _235-340_ **(C)**, and hCDCA7 _264-371_ C339S **(D)** with double stranded DNA oligonucleotides with an unmethylated, hemi-methylated or fully-methylated CpG.

**Fig. S4.**
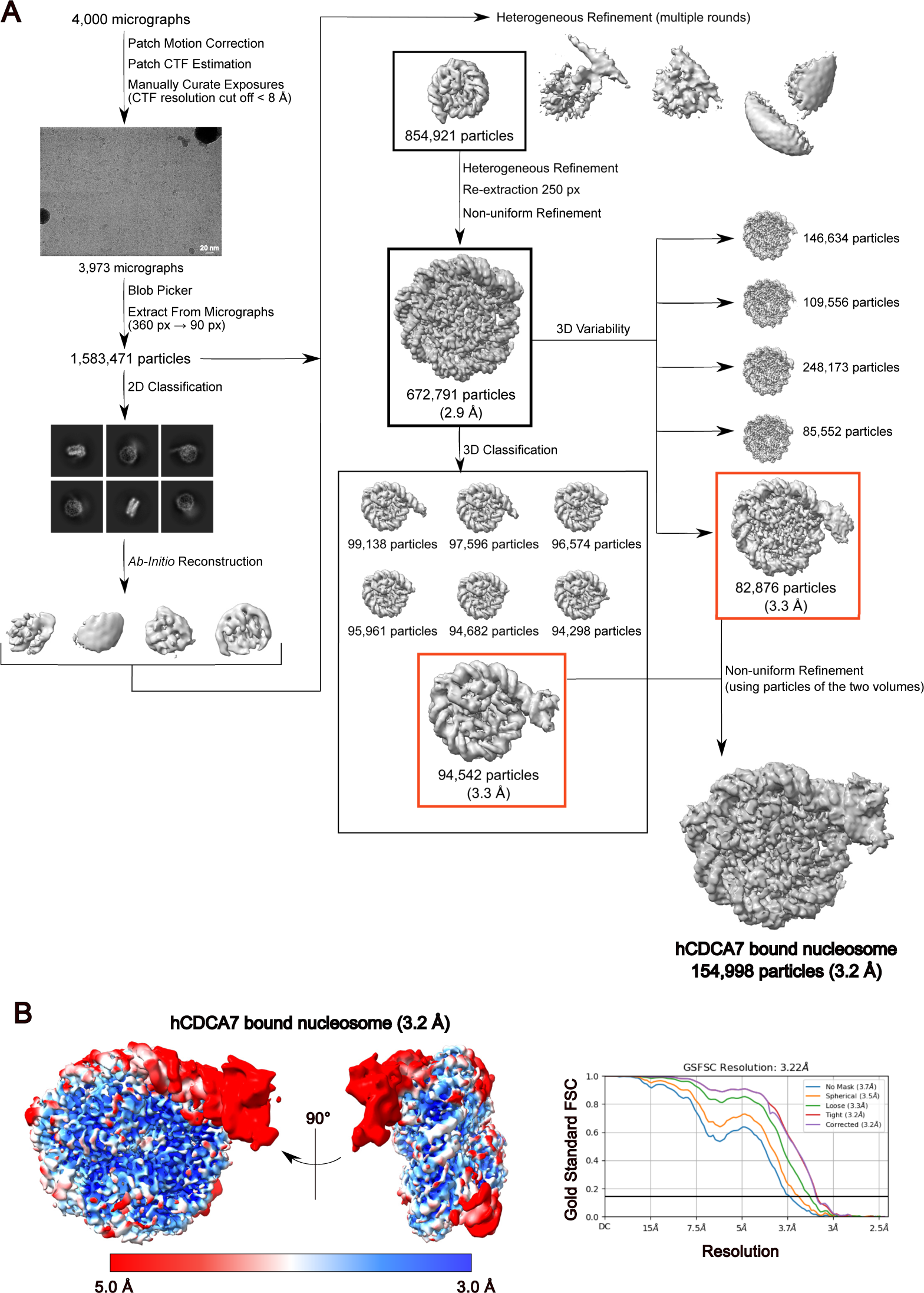
Cryo-EM single particle analysis of hCDCA7 bound to nucleosome. **(A)** Cryo-EM data particle processing and refinement workflow of hCDCA7_264-371_ C339S bound to nucleosome. **(B)** Local resolution of cryo-EM map of hCDCA7_264-371_ C339S bound to nucleosome (left). Fourier shell correlation (FSC) curve of hCDCA7_264-371_ C339S bound to nucleosome (right).

**Fig. S5.**
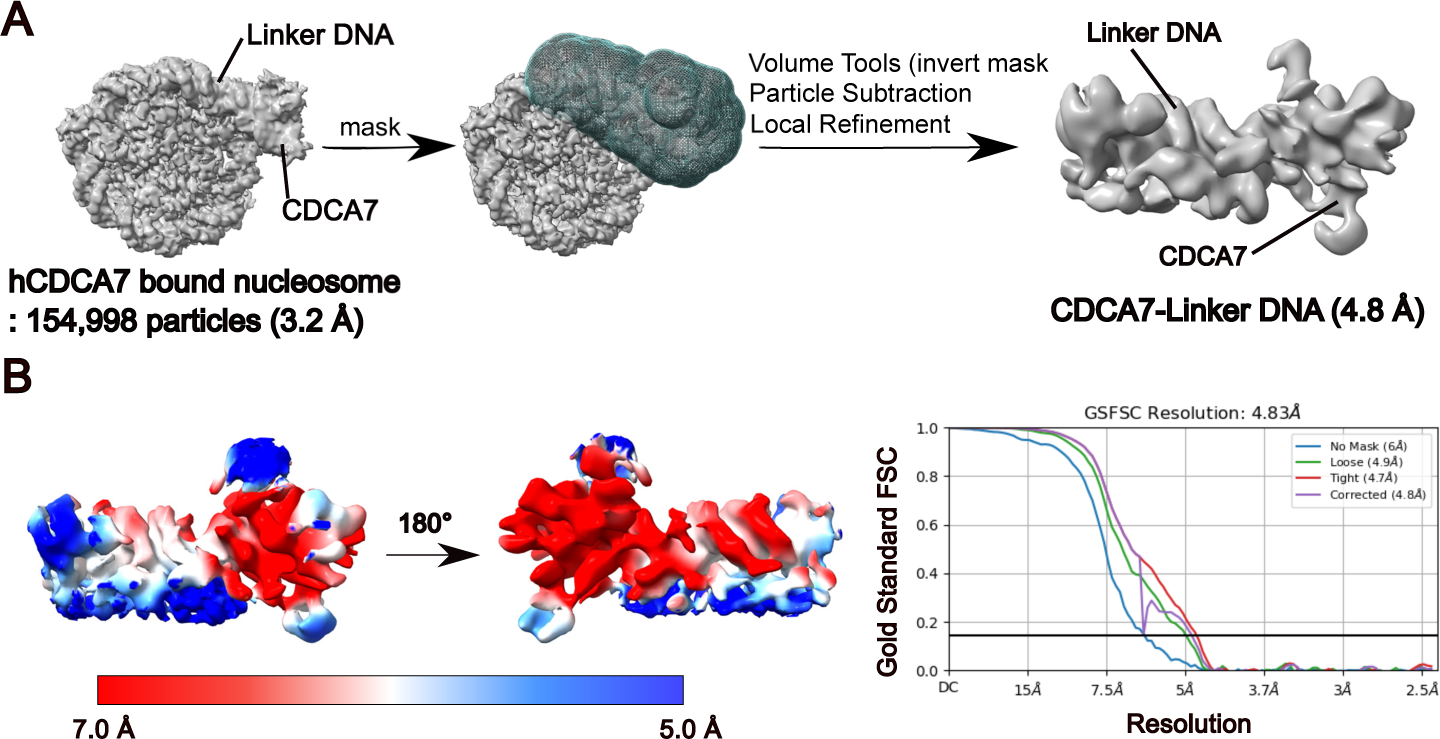
Refinement workflow for cryo-EM map of linker DNA bound to hCDCA7 density. **(A)** Focused refinement of the linker DNA bound by hCDCA7_264-371_ C339S moiety. Left figure shows the cryo-EM map of hCDCA7_264-371_ C339S bound to nucleosome. The mask file is shown as a green mesh (center) covering the hCDCA7_264-371_ C339S moiety bound to the linker DNA. The cryo-EM map corresponding to the hCDCA7_264-371_ C339S moiety bound to the linker DNA was improved by local refinement at 4.83 Å resolution (right). **(B)** Local resolution of the hCDCA7_264-371_ C339S moiety bound to linker DNA. (left). FSC curve of hCDCA7_264-371_ C339S bound to nucleosome (right)

**Fig. S6.**
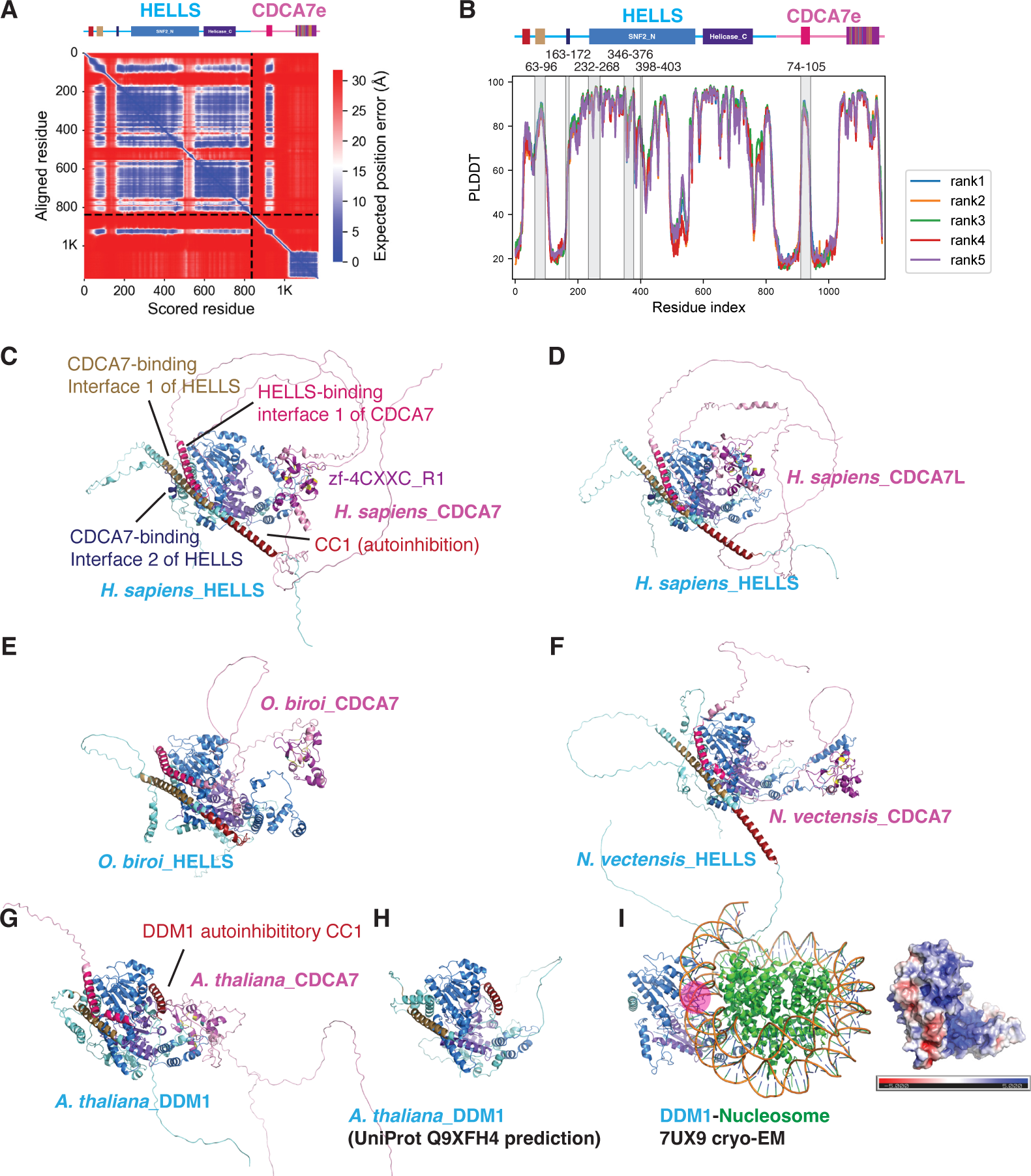
Alphafold2 structure prediction of the CDCA7-HELLS/DDM1 complex. (**A, B**) AlphaFold2 structure prediction analysis of *X. laevis* HELLS and CDCA7e. (**A**) The predicted aligned error map of the best model, with the minimum inter-chain predicted aligned error of 1.7 Å. (**B**) PLDDT scores of the top five predicted models, with the interface highlighted on top of the figure. The top five predictions were converged (region is shaded in gray), and the interface has relatively high PLDDT scores, with the average value of 63. (**C-H**) AlphaFold2 structure prediction of HELLS/DDM1 of indicated species in complex with CDCA7 and CDCA7 paralogs. (**I**) Left; atomic model of DDM1-nucleosome complex cryo-EM structure (7UX9). Right; surface electrostatic potential of DDM1. The DNA-binding positively charged groove, which is predicted to be occupied by the autoinhibitory CC by AlphaFold2 models, is marked with a pink circle.

**Fig. S7.**
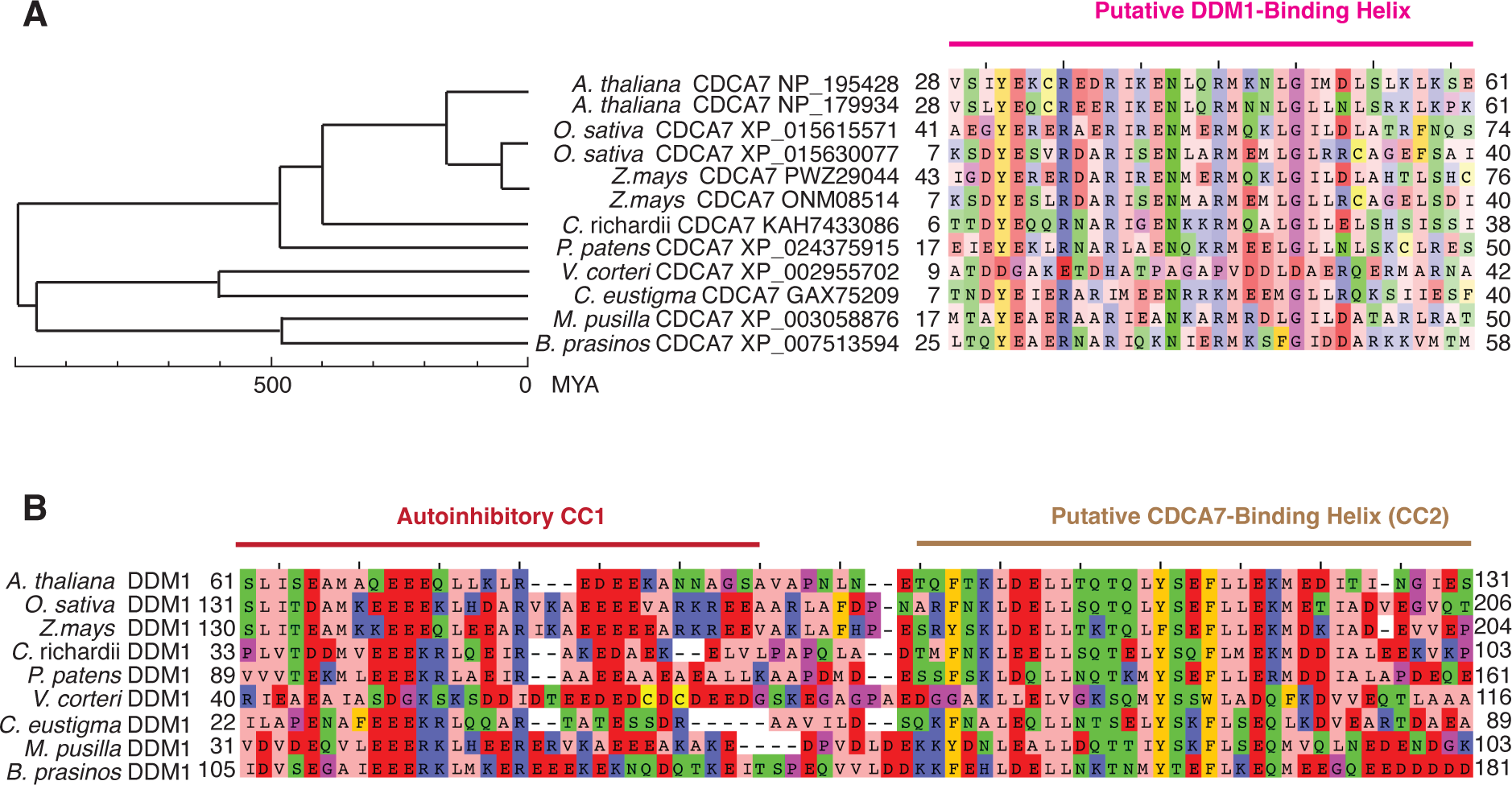
Evolutionary conservation of putative DDM1-CDCA7 interaction interfaces in green plants. **(A)** Sequence alignment of putative DDM1-binding helix of CDCA7 homologs in green plants. *O. sativa*, *Oryza sativa* (rice); *Z. mays*, *Zea mays* (corn); *C. richardii*, *Ceratopteris richardii* (fern); *P. patens*, *Physcomitrium patens* (moss); *V. carteri*, *Volvox carteri* (colonial green alga); *C. eustigma*, *Chlamydomonas eustigma* (unicellular green alga); *M. pusilla*, *Micromonas pusilla* (unicellular green alga); *B. prasinos*, *Bathycoccus prasinos* (marine green alga). (**(B)** Sequence alignment of the putative autoinhibitory CC1 and the CDCA7-binding CC2 of DDM1 homologs in green plants.

**Fig. S8.**
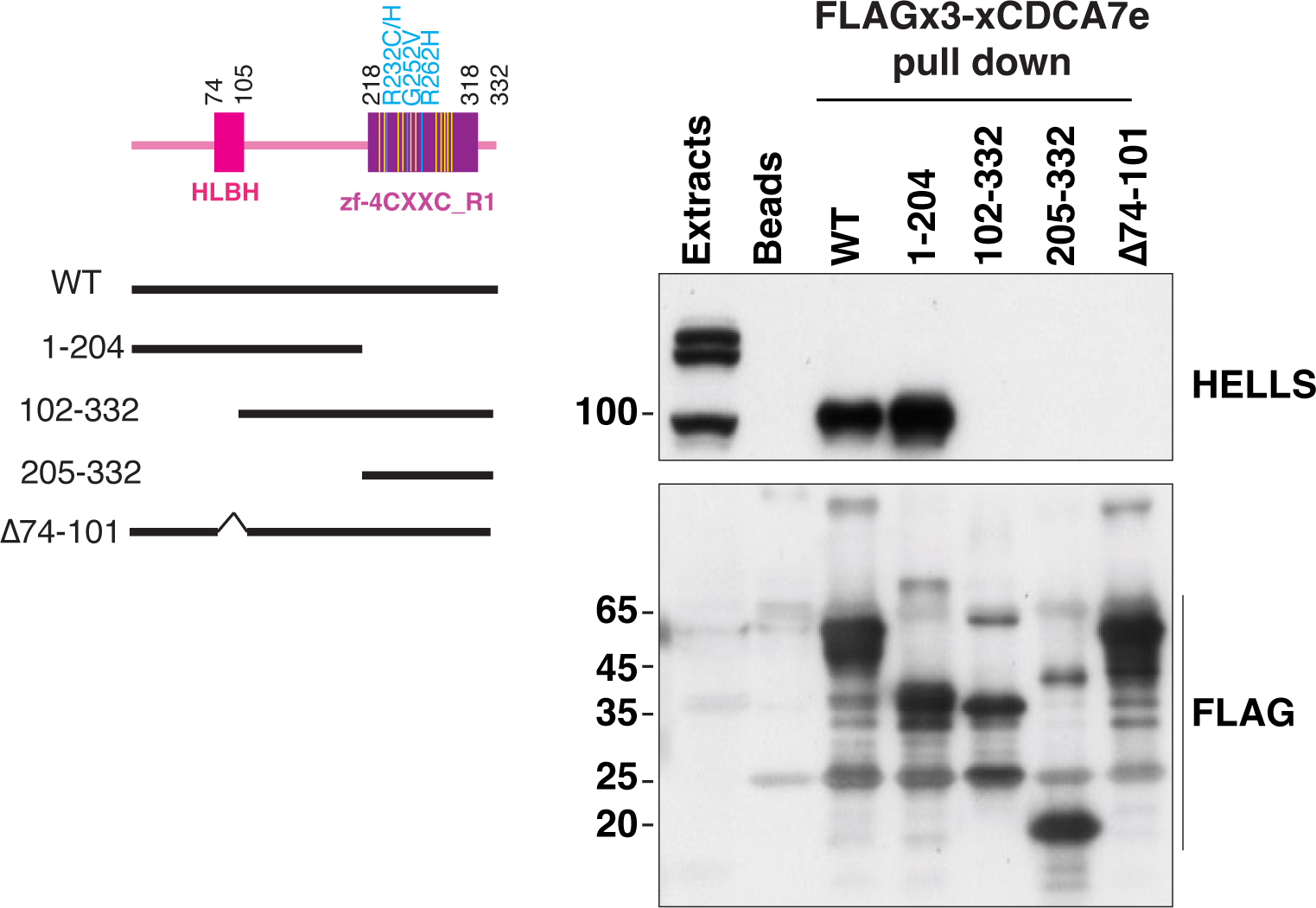
N-terminal CDCA7 segment lacking the zf-4CXXC_R1 domain is sufficient for HELLS binding. Wildtype (WT) or truncated versions of recombinant FLAG3-tagged *X. laevis* CDCA7e proteins were incubated with *Xenopus* egg extracts, followed by immunoprecipitation with anti-FLAG coupled beads. Isolated proteins were analyzed by western blotting using anti-HELLS and anti-FLAG antibodies.

**Fig. S9.**
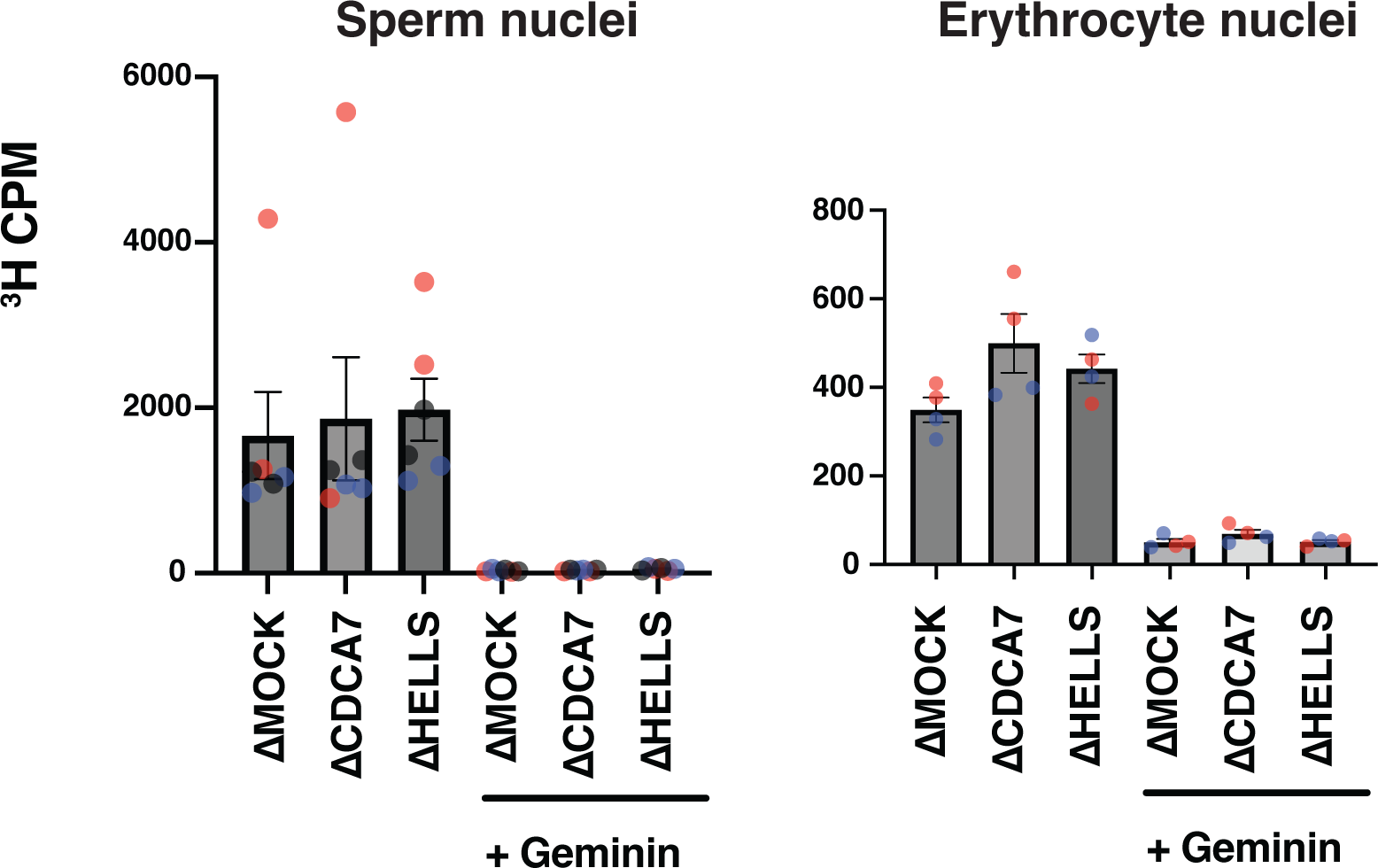
CDCA7 and HELLS are not required for global maintenance DNA methylation in *Xenopus* egg extracts. **(A)** *X. laevis* sperm nuclei **(A)** or erythrocyte nuclei **(B)** were incubated with egg extracts for 60 min with *S*-[methyl-^3^H]-adenosyl-L-methionine with or without geminin, which inhibits DNA replication initiation. Radioactivity associated with chromosomal DNA is measured. Results include three biological replicates **(A)** or two biological replicates **(B)**, each of which includes two technical replicates (shown in the same color). Geminin effectively inhibited DNA incorporation of ^3^H, demonstrating that DNA methylation of sperm chromatin depends on DNA replication.

**Fig. S10.**
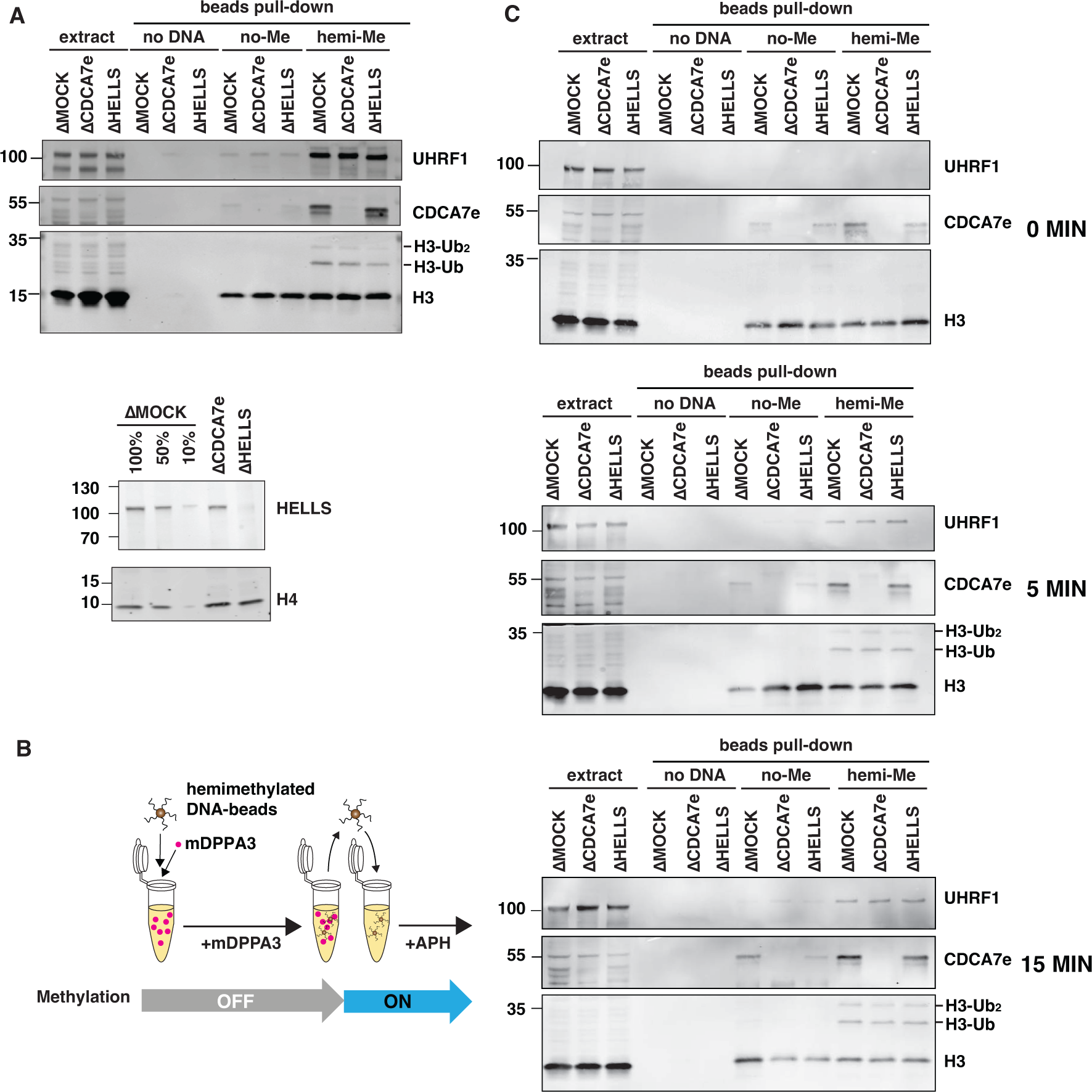
CDCA7e and HELLS are not required for histone H3 ubiquitylation on hemimethylated DNA-beads in *Xenopus* egg extracts. **(A)** Beads coated with unmethylated pBlueScript DNA or hemimethylated pBlueScript DNA were incubated with interphase mock IgG-depleted (ΔMOCK), CDCA7e-depleted (ΔCDCA7e), or HELLS-depleted (ΔHELLS) *Xenopus* egg extracts for 60 min. Beads were collected and analyzed by western blotting. Bottom panel shows effective HELLS depletion. (**B, C**) Beads coated with unmethylated pBlueScript DNA or hemimethylated pBlueScript DNA were incubated with ΔMOCK, ΔCDCA7e, or ΔHELLS egg extracts for 60 min in the presence of 1.3 µM mDPPA3, which inhibits binding of UHRF1 and H3 ubiquitylation. During this preincubation, nucleosomes assemble on DNA beads without DNA methylation. Beads were then transferred to corresponding depleted interphase extracts that contained aphidicolin (APH) but not mDPPA3. After 0-, 5-, or 15-min incubation, beads were collected and analyzed by western blotting.

**Table S1.**
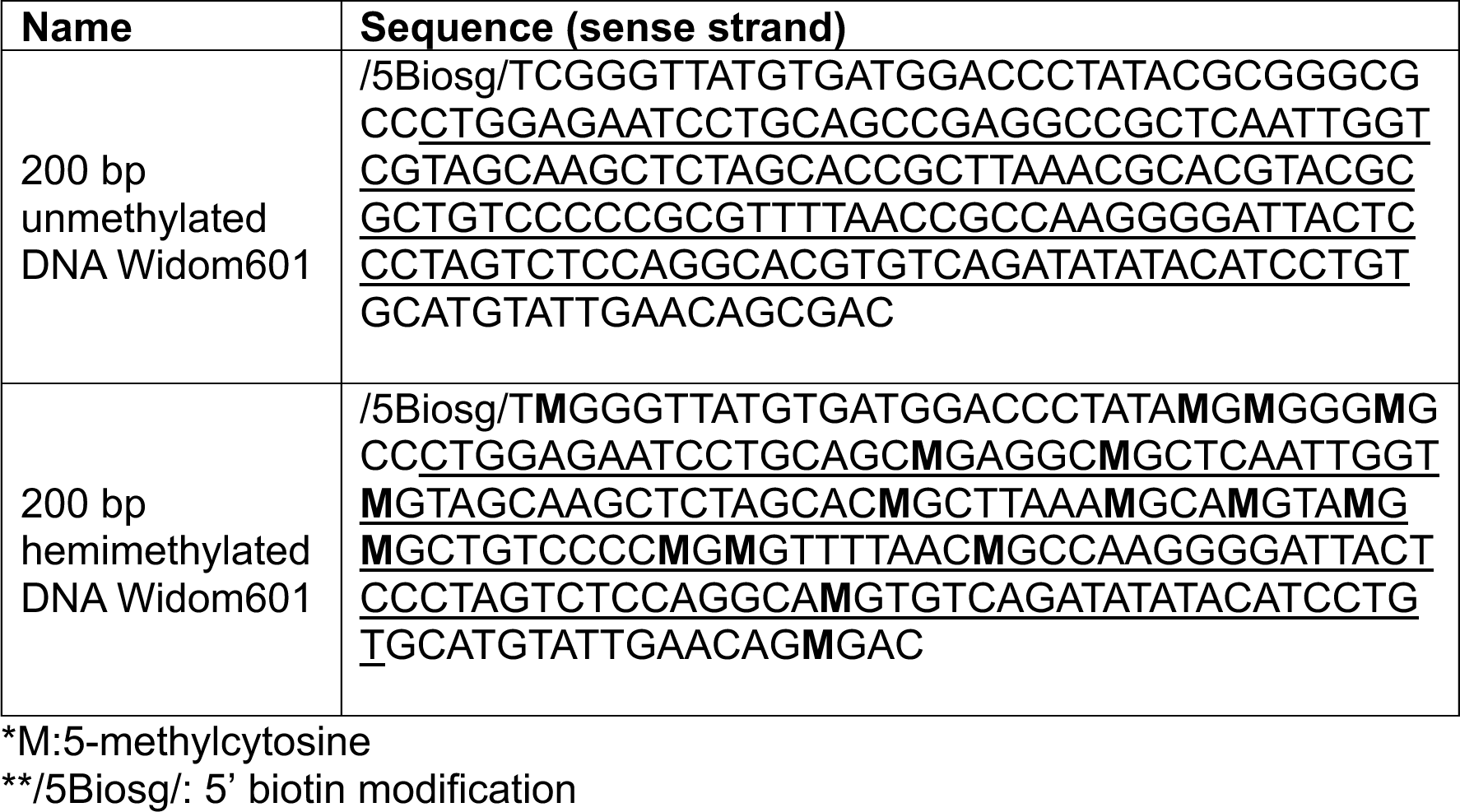
DNA ultramer sequence used for DNA pull-downs.

**Table S2.**
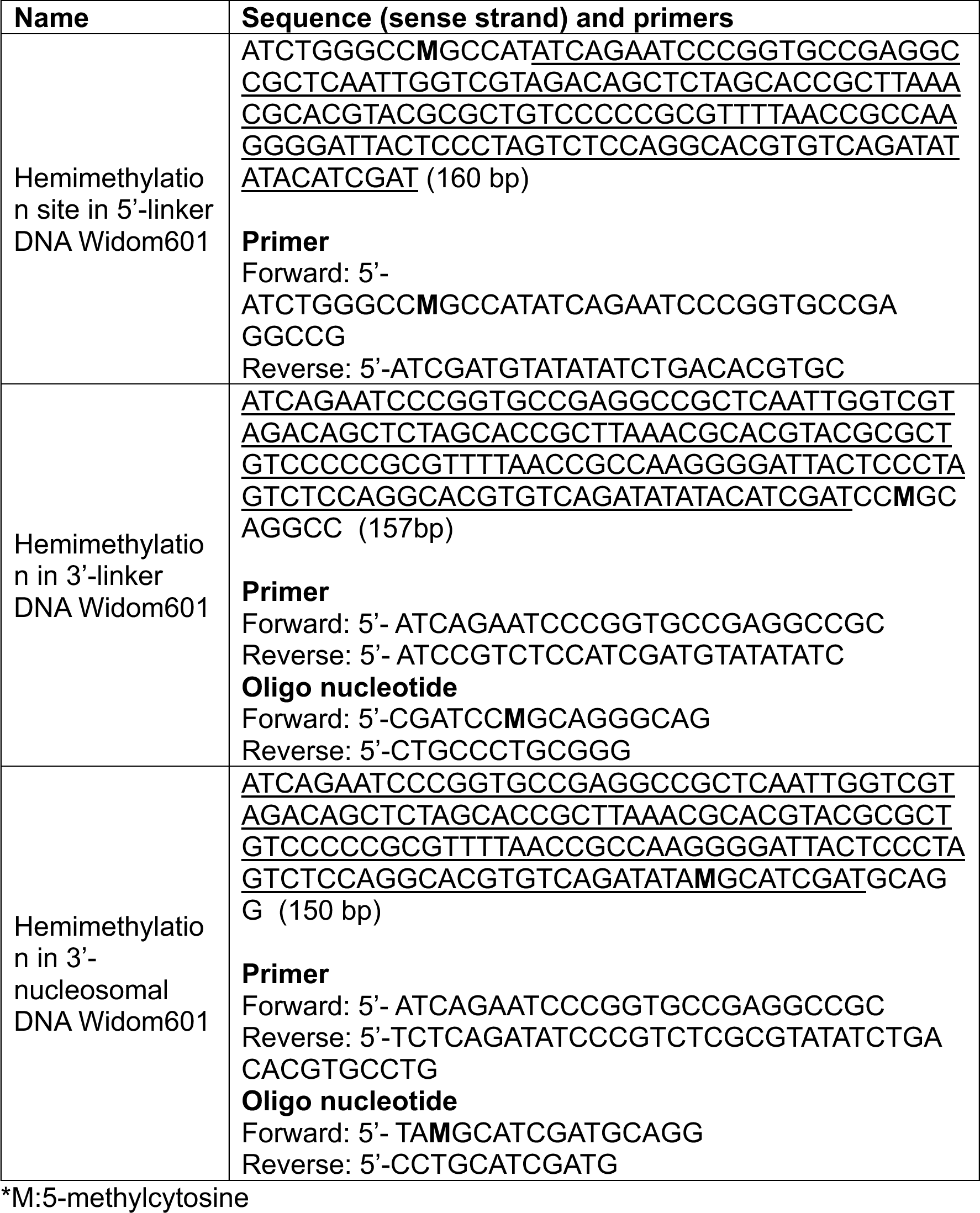
DNA sequence used for nucleosome reconstruction.

**Table S3.** Cryo-EM data collection statics for hCDCA7:nucleosome.

|  | **hCDCA7:nucleosome** | **hCDCA7:linker DNA**<br>**(focused map)** |
| --- | --- | --- |
| EMDB number | EMD-38198 | EMD-38199 |
| Microscope | Krios G4 (RIKEN BDR) |  |
| Voltage (keV) | 300 |  |
| Camera | K3/BioQuantum |  |
| Magnification | 105,000 |  |
| Pixel size at detector (Å) | 0.83 |  |
| Total electron exposure (e<sup>-</sup>/Å<sup>2</sup>) | 60.725 |  |
| Exposure rate (e<sup>-</sup>/pixel/sec) | 18.987 |  |
| Exposure time (sec) | 2.2 |  |
| Defocus range (µm) | 0.6-1.6 (interval: 0.2) |  |
| Number of frames | 48 |  |
| Energy filter slit width | 15 |  |
| Micrographs collected (no.) | 4,000 |  |
| Initial particle images (no.) | 1,652,465 | 672,791 |
| Final particle images (no.) | 154,998 | 154,998 |
| Map resolution (Å) FSC threshold | 3.18 | 4.83 |
| Automation software | EPU |  |

## References

1. O. Deniz, J. M. Frost, M. R. Branco, Regulation of transposable elements by DNA modifications. Nat Rev Genet 20, 417–431 (2019).

2. J. Casadesus, D. Low, Epigenetic gene regulation in the bacterial world. Microbiol Mol Biol Rev 70, 830–856 (2006).

3. T. Dimitriu, M. D. Szczelkun, E. R. Westra, Evolutionary Ecology and Interplay of Prokaryotic Innate and Adaptive Immune Systems. Current biology : CB 30, R1189–R1202 (2020).

4. A. L. Mattei, N. Bailly, A. Meissner, DNA methylation: a historical perspective. Trends Genet 38, 676–707 (2022).

5. E. A. Miska, A. C. Ferguson-Smith, Transgenerational inheritance: Models and mechanisms of non-DNA sequence-based inheritance. Science 354, 59–63 (2016).

6. M. V. C. Greenberg, D. Bourc’his, The diverse roles of DNA methylation in mammalian development and disease. Nat Rev Mol Cell Biol 20, 590–607 (2019).

7. A. Nishiyama, M. Nakanishi, Navigating the DNA methylation landscape of cancer. Trends Genet 37, 1012–1027 (2021).

8. K. D. Robertson, DNA methylation and human disease. Nat Rev Genet 6, 597–610 (2005).

9. R. Lowe, C. Barton, C. A. Jenkins, C. Ernst, O. Forman, D. S. Fernandez-Twinn, C. Bock, S. J. Rossiter, C. G. Faulkes, S. E. Ozanne, L. Walter, D. T. Odom, C. Mellersh, V. K. Rakyan, Ageing-associated DNA methylation dynamics are a molecular readout of lifespan variation among mammalian species. Genome Biol 19, 22 (2018).

10. M. Ehrlich, K. Jackson, C. Weemaes, Immunodeficiency, centromeric region instability, facial anomalies syndrome (ICF). Orphanet J Rare Dis 1, 2 (2006).

11. M. Vukic, L. Daxinger, DNA methylation in disease: Immunodeficiency, Centromeric instability, Facial anomalies syndrome. Essays Biochem 63, 773–783 (2019).

12. R. S. Hansen, C. Wijmenga, P. Luo, A. M. Stanek, T. K. Canfield, C. M. Weemaes, S. M. Gartler, The DNMT3B DNA methyltransferase gene is mutated in the ICF immunodeficiency syndrome. Proc Natl Acad Sci U S A 96, 14412–14417 (1999).

13. M. Okano, D. W. Bell, D. A. Haber, E. Li, DNA methyltransferases Dnmt3a and Dnmt3b are essential for de novo methylation and mammalian development. Cell 99, 247–257 (1999).

14. J. C. de Greef, J. Wang, J. Balog, J. T. den Dunnen, R. R. Frants, K. R. Straasheijm, C. Aytekin, M. van der Burg, L. Duprez, A. Ferster, A. R. Gennery, G. Gimelli, I. Reisli, C. Schuetz, A. Schulz, D. Smeets, Y. Sznajer, C. Wijmenga, M. C. van Eggermond, M. M. van Ostaijen-Ten Dam, A. C. Lankester, M. J. D. van Tol, P. J. van den Elsen, C. M. Weemaes, S. M. van der Maarel, Mutations in ZBTB24 are associated with immunodeficiency, centromeric instability, and facial anomalies syndrome type 2. Am J Hum Genet 88, 796–804 (2011).

15. P. E. Thijssen, Y. Ito, G. Grillo, J. Wang, G. Velasco, H. Nitta, M. Unoki, M. Yoshihara, M. Suyama, Y. Sun, R. J. Lemmers, J. C. de Greef, A. Gennery, P. Picco, B. Kloeckener-Gruissem, T. Gungor, I. Reisli, C. Picard, K. Kebaili, B. Roquelaure, T. Iwai, I. Kondo, T. Kubota, M. M. van Ostaijen-Ten Dam, M. J. van Tol, C. Weemaes, C. Francastel, S. M. van der Maarel, H. Sasaki, Mutations in CDCA7 and HELLS cause immunodeficiency-centromeric instability-facial anomalies syndrome. Nature communications 6, 7870 (2015).

16. M. Unoki, Chromatin remodeling in replication-uncoupled maintenance DNA methylation and chromosome stability: Insights from ICF syndrome studies. Genes Cells 26, 349–359 (2021).

17. M. Unoki, G. Velasco, S. Kori, K. Arita, Y. Daigaku, W. K. A. Yeung, A. Fujimoto, H. Ohashi, T. Kubota, K. Miyake, H. Sasaki, Novel compound heterozygous mutations in UHRF1 are associated with atypical immunodeficiency, centromeric instability and facial anomalies syndrome with distinctive genome-wide DNA hypomethylation. Hum Mol Genet 32, 1439–1456 (2023).

18. D. S. Dunican, H. A. Cruickshanks, M. Suzuki, C. A. Semple, T. Davey, R. J. Arceci, J. Greally, I. R. Adams, R. R. Meehan, Lsh regulates LTR retrotransposon repression independently of Dnmt3b function. Genome Biol 14, R146 (2013).

19. D. S. Dunican, S. Pennings, R. R. Meehan, Lsh Is Essential for Maintaining Global DNA Methylation Levels in Amphibia and Fish and Interacts Directly with Dnmt1. Biomed Res Int 2015, 740637 (2015).

20. K. Myant, A. Termanis, A. Y. Sundaram, T. Boe, C. Li, C. Merusi, J. Burrage, J. I. de Las Heras, I. Stancheva, LSH and G9a/GLP complex are required for developmentally programmed DNA methylation. Genome Res 21, 83–94 (2011).

21. W. Yu, C. McIntosh, R. Lister, I. Zhu, Y. Han, J. Ren, D. Landsman, E. Lee, V. Briones, M. Terashima, R. Leighty, J. R. Ecker, K. Muegge, Genome-wide DNA methylation patterns in LSH mutant reveals de-repression of repeat elements and redundant epigenetic silencing pathways. Genome Res 24, 1613–1623 (2014).

22. A. Vongs, T. Kakutani, R. A. Martienssen, E. J. Richards, Arabidopsis thaliana DNA methylation mutants. Science 260, 1926–1928 (1993).

23. A. Miura, S. Yonebayashi, K. Watanabe, T. Toyama, H. Shimada, T. Kakutani, Mobilization of transposons by a mutation abolishing full DNA methylation in Arabidopsis. Nature 411, 212–214 (2001).

24. K. Dennis, T. Fan, T. Geiman, Q. Yan, K. Muegge, Lsh, a member of the SNF2 family, is required for genome-wide methylation. Genes Dev 15, 2940–2944 (2001).

25. M. Han, J. Li, Y. Cao, Y. Huang, W. Li, H. Zhu, Q. Zhao, J. J. Han, Q. Wu, J. Li, J. Feng, J. Wong, A role for LSH in facilitating DNA methylation by DNMT1 through enhancing UHRF1 chromatin association. Nucleic Acids Res 48, 12116–12134 (2020).

26. D. B. Lyons, D. Zilberman, DDM1 and Lsh remodelers allow methylation of DNA wrapped in nucleosomes. Elife 6, (2017).

27. C. Jenness, S. Giunta, M. M. Muller, H. Kimura, T. W. Muir, H. Funabiki, HELLS and CDCA7 comprise a bipartite nucleosome remodeling complex defective in ICF syndrome. Proc Natl Acad Sci U S A 115, E876–E885 (2018).

28. H. Funabiki, I. E. Wassing, Q. Jia, J. D. Luo, T. Carroll, Coevolution of the CDCA7-HELLS ICF-related nucleosome remodeling complex and DNA methyltransferases. Elife 12, (2023).

29. F. Lyko, The DNA methyltransferase family: a versatile toolkit for epigenetic regulation. Nat Rev Genet 19, 81–92 (2018).

30. K. Arita, M. Ariyoshi, H. Tochio, Y. Nakamura, M. Shirakawa, Recognition of hemi-methylated DNA by the SRA protein UHRF1 by a base-flipping mechanism. Nature 455, 818–821 (2008).

31. G. V. Avvakumov, J. R. Walker, S. Xue, Y. Li, S. Duan, C. Bronner, C. H. Arrowsmith, S. Dhe-Paganon, Structural basis for recognition of hemi-methylated DNA by the SRA domain of human UHRF1. Nature 455, 822–825 (2008).

32. H. Hashimoto, J. R. Horton, X. Zhang, M. Bostick, S. E. Jacobsen, X. Cheng, The SRA domain of UHRF1 flips 5-methylcytosine out of the DNA helix. Nature 455, 826–829 (2008).

33. M. Bostick, J. K. Kim, P. O. Esteve, A. Clark, S. Pradhan, S. E. Jacobsen, UHRF1 plays a role in maintaining DNA methylation in mammalian cells. Science 317, 1760–1764 (2007).

34. J. Sharif, M. Muto, S. Takebayashi, I. Suetake, A. Iwamatsu, T. A. Endo, J. Shinga, Y. Mizutani-Koseki, T. Toyoda, K. Okamura, S. Tajima, K. Mitsuya, M. Okano, H. Koseki, The SRA protein Np95 mediates epigenetic inheritance by recruiting Dnmt1 to methylated DNA. Nature 450, 908–912 (2007).

35. A. Nishiyama, L. Yamaguchi, J. Sharif, Y. Johmura, T. Kawamura, K. Nakanishi, S. Shimamura, K. Arita, T. Kodama, F. Ishikawa, H. Koseki, M. Nakanishi, Uhrf1-dependent H3K23 ubiquitylation couples maintenance DNA methylation and replication. Nature 502, 249–253 (2013).

36. M. Mancini, E. Magnani, F. Macchi, I. M. Bonapace, The multi-functionality of UHRF1: epigenome maintenance and preservation of genome integrity. Nucleic Acids Res 49, 6053–6068 (2021).

37. A. Nishiyama, C. B. Mulholland, S. Bultmann, S. Kori, A. Endo, Y. Saeki, W. Qin, C. Trummer, Y. Chiba, H. Yokoyama, S. Kumamoto, T. Kawakami, H. Hojo, G. Nagae, H. Aburatani, K. Tanaka, K. Arita, H. Leonhardt, M. Nakanishi, Two distinct modes of DNMT1 recruitment ensure stable maintenance DNA methylation. Nature communications 11, 1222 (2020).

38. S. Ishiyama, A. Nishiyama, Y. Saeki, K. Moritsugu, D. Morimoto, L. Yamaguchi, N. Arai, R. Matsumura, T. Kawakami, Y. Mishima, H. Hojo, S. Shimamura, F. Ishikawa, S. Tajima, K. Tanaka, M. Ariyoshi, M. Shirakawa, M. Ikeguchi, A. Kidera, I. Suetake, K. Arita, M. Nakanishi, Structure of the Dnmt1 Reader Module Complexed with a Unique Two-Mono-Ubiquitin Mark on Histone H3 Reveals the Basis for DNA Methylation Maintenance. Mol Cell 68, 350–360 e357 (2017).

39. X. Ming, Z. Zhang, Z. Zou, C. Lv, Q. Dong, Q. He, Y. Yi, Y. Li, H. Wang, B. Zhu, Kinetics and mechanisms of mitotic inheritance of DNA methylation and their roles in aging-associated methylome deterioration. Cell Res 30, 980–996 (2020).

40. C. Zierhut, C. Jenness, H. Kimura, H. Funabiki, Nucleosomal regulation of chromatin composition and nuclear assembly revealed by histone depletion. Nat Struct Mol Biol 21, 617–625 (2014).

41. M. Felle, H. Hoffmeister, J. Rothammer, A. Fuchs, J. H. Exler, G. Langst, Nucleosomes protect DNA from DNA methylation in vivo and in vitro. Nucleic Acids Res 39, 6956–6969 (2011).

42. M. Okuwaki, A. Verreault, Maintenance DNA methylation of nucleosome core particles. The Journal of biological chemistry 279, 2904–2912 (2004).

43. A. K. Robertson, T. M. Geiman, U. T. Sankpal, G. L. Hager, K. D. Robertson, Effects of chromatin structure on the enzymatic and DNA binding functions of DNA methyltransferases DNMT1 and Dnmt3a in vitro. Biochem Biophys Res Commun 322, 110–118 (2004).

44. H. Takeshima, I. Suetake, H. Shimahara, K. Ura, S. Tate, S. Tajima, Distinct DNA methylation activity of Dnmt3a and Dnmt3b towards naked and nucleosomal DNA. J Biochem 139, 503–515 (2006).

45. A. Schrader, T. Gross, V. Thalhammer, G. Langst, Characterization of Dnmt1 Binding and DNA Methylation on Nucleosomes and Nucleosomal Arrays. PloS one 10, e0140076 (2015).

46. Q. Zhao, J. Zhang, R. Chen, L. Wang, B. Li, H. Cheng, X. Duan, H. Zhu, W. Wei, J. Li, Q. Wu, J. D. Han, W. Yu, S. Gao, G. Li, J. Wong, Dissecting the precise role of H3K9 methylation in crosstalk with DNA maintenance methylation in mammals. Nature communications 7, 12464 (2016).

47. J. Brzeski, A. Jerzmanowski, Deficient in DNA methylation 1 (DDM1) defines a novel family of chromatin-remodeling factors. The Journal of biological chemistry 278, 823–828 (2003).

48. S. C. Lee, D. W. Adams, J. J. Ipsaro, J. Cahn, J. Lynn, H. S. Kim, B. Berube, V. Major, J. P. Calarco, C. LeBlanc, S. Bhattacharjee, U. Ramu, D. Grimanelli, Y. Jacob, P. Voigt, L. Joshua-Tor, R. A. Martienssen, Chromatin remodeling of histone H3 variants by DDM1 underlies epigenetic inheritance of DNA methylation. Cell 186, 4100–4116 e4115 (2023).

49. M. Unoki, H. Funabiki, G. Velasco, C. Francastel, H. Sasaki, CDCA7 and HELLS mutations undermine nonhomologous end joining in centromeric instability syndrome. J Clin Invest 129, 78–92 (2019).

50. C. E. Shamu, A. W. Murray, Sister chromatid separation in frog egg extracts requires DNA topoisomerase II activity during anaphase. The Journal of cell biology 117, 921–934 (1992).

51. C. B. Mulholland, A. Nishiyama, J. Ryan, R. Nakamura, M. Yigit, I. M. Gluck, C. Trummer, W. Qin, M. D. Bartoschek, F. R. Traube, E. Parsa, E. Ugur, M. Modic, A. Acharya, P. Stolz, C. Ziegenhain, M. Wierer, W. Enard, T. Carell, D. C. Lamb, H. Takeda, M. Nakanishi, S. Bultmann, H. Leonhardt, Recent evolution of a TET-controlled and DPPA3/STELLA-driven pathway of passive DNA demethylation in mammals. Nature communications 11, 5972 (2020).

52. K. Hata, N. Kobayashi, K. Sugimura, W. Qin, D. Haxholli, Y. Chiba, S. Yoshimi, G. Hayashi, H. Onoda, T. Ikegami, C. B. Mulholland, A. Nishiyama, M. Nakanishi, H. Leonhardt, T. Konuma, K. Arita, Structural basis for the unique multifaceted interaction of DPPA3 with the UHRF1 PHD finger. Nucleic Acids Res 50, 12527–12542 (2022).

53. Y. Li, Z. Zhang, J. Chen, W. Liu, W. Lai, B. Liu, X. Li, L. Liu, S. Xu, Q. Dong, M. Wang, X. Duan, J. Tan, Y. Zheng, P. Zhang, G. Fan, J. Wong, G. L. Xu, Z. Wang, H. Wang, S. Gao, B. Zhu, Stella safeguards the oocyte methylome by preventing de novo methylation mediated by DNMT1. Nature 564, 136–140 (2018).

54. J. A. Wohlschlegel, B. T. Dwyer, S. K. Dhar, C. Cvetic, J. C. Walter, A. Dutta, Inhibition of eukaryotic DNA replication by geminin binding to Cdt1. Science 290, 2309–2312 (2000).

55. J. Jumper, R. Evans, A. Pritzel, T. Green, M. Figurnov, O. Ronneberger, K. Tunyasuvunakool, R. Bates, A. Zidek, A. Potapenko, A. Bridgland, C. Meyer, S. A. A. Kohl, A. J. Ballard, A. Cowie, B. Romera-Paredes, S. Nikolov, R. Jain, J. Adler, T. Back, S. Petersen, D. Reiman, E. Clancy, M. Zielinski, M. Steinegger, M. Pacholska, T. Berghammer, S. Bodenstein, D. Silver, O. Vinyals, A. W. Senior, K. Kavukcuoglu, P. Kohli, D. Hassabis, Highly accurate protein structure prediction with AlphaFold. Nature 596, 583–589 (2021).

56. M. Mirdita, K. Schutze, Y. Moriwaki, L. Heo, S. Ovchinnikov, M. Steinegger, ColabFold: making protein folding accessible to all. Nat Methods 19, 679–682 (2022).

57. W. Nartey, A. A. Goodarzi, G. J. Williams, Cryo-EM structure of DDM1-HELLS chimera bound to nucleosome reveals a mechanism of chromatin remodeling and disease regulation. bioRxiv, 2023.2008.2009.551721 (2023).

58. G. Velasco, G. Grillo, N. Touleimat, L. Ferry, I. Ivkovic, F. Ribierre, J. F. Deleuze, S. Chantalat, C. Picard, C. Francastel, Comparative methylome analysis of ICF patients identifies heterochromatin loci that require ZBTB24, CDCA7 and HELLS for their methylated state. Hum Mol Genet, (2018).

59. A. Zemach, M. Y. Kim, P. H. Hsieh, D. Coleman-Derr, L. Eshed-Williams, K. Thao, S. L. Harmer, D. Zilberman, The Arabidopsis nucleosome remodeler DDM1 allows DNA methyltransferases to access H1-containing heterochromatin. Cell 153, 193–205 (2013).

60. H. Tagami, D. Ray-Gallet, G. Almouzni, Y. Nakatani, Histone H3.1 and H3.3 complexes mediate nucleosome assembly pathways dependent or independent of DNA synthesis. Cell 116, 51–61 (2004).

61. G. Almouzni, M. Mechali, Assembly of spaced chromatin promoted by DNA synthesis in extracts from Xenopus eggs. EMBO J 7, 665–672 (1988).

62. J. Burrage, A. Termanis, A. Geissner, K. Myant, K. Gordon, I. Stancheva, The SNF2 family ATPase LSH promotes phosphorylation of H2AX and efficient repair of DNA double-strand breaks in mammalian cells. J Cell Sci 125, 5524–5534 (2012).

63. K. Luger, A. W. Mader, R. K. Richmond, D. F. Sargent, T. J. Richmond, Crystal structure of the nucleosome core particle at 2.8 A resolution. Nature 389, 251–260 (1997).

64. J. Song, M. Teplova, S. Ishibe-Murakami, D. J. Patel, Structure-based mechanistic insights into DNMT1-mediated maintenance DNA methylation. Science 335, 709–712 (2012).

65. H. S. Rhee, A. R. Bataille, L. Zhang, B. F. Pugh, Subnucleosomal structures and nucleosome asymmetry across a genome. Cell 159, 1377–1388 (2014).

66. A. Stutzer, S. Liokatis, A. Kiesel, D. Schwarzer, R. Sprangers, J. Soding, P. Selenko, W. Fischle, Modulations of DNA Contacts by Linker Histones and Post-translational Modifications Determine the Mobility and Modifiability of Nucleosomal H3 Tails. Mol Cell 61, 247–259 (2016).

67. D. Angelov, J. M. Vitolo, V. Mutskov, S. Dimitrov, J. J. Hayes, Preferential interaction of the core histone tail domains with linker DNA. Proc Natl Acad Sci U S A 98, 6599–6604 (2001).

68. Y. Peng, S. Li, A. Onufriev, D. Landsman, A. R. Panchenko, Binding of regulatory proteins to nucleosomes is modulated by dynamic histone tails. Nature communications 12, 5280 (2021).

69. B. M. Foster, P. Stolz, C. B. Mulholland, A. Montoya, H. Kramer, S. Bultmann, T. Bartke, Critical Role of the UBL Domain in Stimulating the E3 Ubiquitin Ligase Activity of UHRF1 toward Chromatin. Mol Cell 72, 739–752 e739 (2018).

70. R. M. Vaughan, B. M. Dickson, M. F. Whelihan, A. L. Johnstone, E. M. Cornett, M. A. Cheek, C. A. Ausherman, M. W. Cowles, Z. W. Sun, S. B. Rothbart, Chromatin structure and its chemical modifications regulate the ubiquitin ligase substrate selectivity of UHRF1. Proc Natl Acad Sci U S A 115, 8775–8780 (2018).

71. X. Liu, Q. Gao, P. Li, Q. Zhao, J. Zhang, J. Li, H. Koseki, J. Wong, UHRF1 targets DNMT1 for DNA methylation through cooperative binding of hemi-methylated DNA and methylated H3K9. Nature communications 4, 1563 (2013).

72. A. Zemach, I. E. McDaniel, P. Silva, D. Zilberman, Genome-wide evolutionary analysis of eukaryotic DNA methylation. Science 328, 916–919 (2010).

73. S. Feng, S. J. Cokus, X. Zhang, P. Y. Chen, M. Bostick, M. G. Goll, J. Hetzel, J. Jain, S. H. Strauss, M. E. Halpern, C. Ukomadu, K. C. Sadler, S. Pradhan, M. Pellegrini, S. E. Jacobsen, Conservation and divergence of methylation patterning in plants and animals. Proc Natl Acad Sci U S A 107, 8689–8694 (2010).

74. R. Libbrecht, P. R. Oxley, L. Keller, D. J. Kronauer, Robust DNA Methylation in the Clonal Raider Ant Brain. Current biology : CB 26, 391–395 (2016).

75. I. Ivasyk, L. Olivos-Cisneros, S. Valdes-Rodriguez, M. Droual, H. Jang, R. J. Schmitz, D. J. C. Kronauer, DNMT1 mutant ants develop normally but have disrupted oogenesis. Nature communications 14, 2201 (2023).

76. Y. He, J. Ren, X. Xu, K. Ni, A. Schwader, R. Finney, C. Wang, L. Sun, K. Klarmann, J. Keller, A. Tubbs, A. Nussenzweig, K. Muegge, Lsh/HELLS is required for B lymphocyte development and immunoglobulin class switch recombination. Proc Natl Acad Sci U S A 117, 20100–20108 (2020).

77. C. Spruce, S. Dlamini, G. Ananda, N. Bronkema, H. Tian, K. Paigen, G. W. Carter, C. L. Baker, HELLS and PRDM9 form a pioneer complex to open chromatin at meiotic recombination hot spots. Genes Dev 34, 398–412 (2020).

78. G. Kollarovic, C. E. Topping, E. P. Shaw, A. L. Chambers, The human HELLS chromatin remodelling protein promotes end resection to facilitate homologous recombination and contributes to DSB repair within heterochromatin. Nucleic Acids Res 48, 1872–1885 (2020).

79. M. Unoki, J. Sharif, Y. Saito, G. Velasco, C. Francastel, H. Koseki, H. Sasaki, CDCA7 and HELLS suppress DNA:RNA hybrid-associated DNA damage at pericentromeric repeats. Sci Rep 10, 17865 (2020).

80. X. Xu, K. Ni, Y. He, J. Ren, C. Sun, Y. Liu, M. I. Aladjem, S. Burkett, R. Finney, X. Ding, S. K. Sharan, K. Muegge, The epigenetic regulator LSH maintains fork protection and genomic stability via MacroH2A deposition and RAD51 filament formation. Nature communications 12, 3520 (2021).

81. J. Zhou, X. Lei, S. Shafiq, W. Zhang, Q. Li, K. Li, J. Zhu, Z. Dong, X. J. He, Q. Sun, DDM1-mediated R-loop resolution and H2A.Z exclusion facilitates heterochromatin formation in Arabidopsis. Sci Adv 9, eadg2699 (2023).

82. A. Osakabe, B. Jamge, E. Axelsson, S. A. Montgomery, S. Akimcheva, A. L. Kuehn, R. Pisupati, Z. J. Lorkovic, R. Yelagandula, T. Kakutani, F. Berger, The chromatin remodeler DDM1 prevents transposon mobility through deposition of histone variant H2A.W. Nat Cell Biol 23, 391–400 (2021).

83. S. Kumamoto, A. Nishiyama, Y. Chiba, R. Miyashita, C. Konishi, Y. Azuma, M. Nakanishi, HPF1-dependent PARP activation promotes LIG3-XRCC1-mediated backup pathway of Okazaki fragment ligation. Nucleic Acids Res 49, 5003–5016 (2021).

84. P. T. Lowary, J. Widom, New DNA sequence rules for high affinity binding to histone octamer and sequence-directed nucleosome positioning. J Mol Biol 276, 19–42 (1998).

85. H. Funabiki, A. W. Murray, The Xenopus chromokinesin Xkid is essential for metaphase chromosome alignment and must be degraded to allow anaphase chromosome movement. Cell 102, 411–424 (2000).

86. R. Lebofsky, T. Takahashi, J. C. Walter, DNA replication in nucleus-free Xenopus egg extracts. Methods Mol Biol 521, 229–252 (2009).

87. S. E. Humphries, D. Young, D. Carroll, Chromatin structure of the 5S ribonucleic acid genes of Xenopus laevis. Biochemistry 18, 3223–3231 (1979).

88. K. Arita, S. Isogai, T. Oda, M. Unoki, K. Sugita, N. Sekiyama, K. Kuwata, R. Hamamoto, H. Tochio, M. Sato, M. Ariyoshi, M. Shirakawa, Recognition of modification status on a histone H3 tail by linked histone reader modules of the epigenetic regulator UHRF1. Proc Natl Acad Sci U S A 109, 12950–12955 (2012).

89. K. Mayanagi, K. Saikusa, N. Miyazaki, S. Akashi, K. Iwasaki, Y. Nishimura, K. Morikawa, Y. Tsunaka, Structural visualization of key steps in nucleosome reorganization by human FACT. Sci Rep 9, 10183 (2019).

90. A. Punjani, J. L. Rubinstein, D. J. Fleet, M. A. Brubaker, cryoSPARC: algorithms for rapid unsupervised cryo-EM structure determination. Nat Methods 14, 290–296 (2017).

91. P. Emsley, B. Lohkamp, W. G. Scott, K. Cowtan, Features and development of Coot. Acta Crystallogr D Biol Crystallogr 66, 486–501 (2010).

92. P. V. Afonine, R. W. Grosse-Kunstleve, N. Echols, J. J. Headd, N. W. Moriarty, M. Mustyakimov, T. C. Terwilliger, A. Urzhumtsev, P. H. Zwart, P. D. Adams, Towards automated crystallographic structure refinement with phenix.refine. Acta Crystallogr D Biol Crystallogr 68, 352–367 (2012).

93. R. Evans, M. O’Neill, A. Pritzel, N. Antropova, A. Senior, T. Green, A. Žídek, R. Bates, S. Blackwell, J. Yim, O. Ronneberger, S. Bodenstein, M. Zielinski, A. Bridgland, A. Potapenko, A. Cowie, K. Tunyasuvunakool, R. Jain, E. Clancy, P. Kohli, J. Jumper, D. Hassabis, Protein complex prediction with AlphaFold-Multimer. bioRxiv, 2021.2010.2004.463034 (2022).

94. P. Bryant, G. Pozzati, A. Elofsson, Improved prediction of protein-protein interactions using AlphaFold2. Nature communications 13, 1265 (2022).

95. K. Katoh, K. Misawa, K. Kuma, T. Miyata, MAFFT: a novel method for rapid multiple sequence alignment based on fast Fourier transform. Nucleic Acids Res 30, 3059–3066 (2002).

96. N. P. Brown, C. Leroy, C. Sander, MView: a web-compatible database search or multiple alignment viewer. Bioinformatics 14, 380–381 (1998).

